# Perphenazine-macrocycle conjugates rapidly sequester the Aβ42 monomer and inhibit amyloid formation

**DOI:** 10.1101/2020.11.16.384248

**Authors:** Sarah R. Ball, Julius S. P Adamson, Michael A. Sullivan, Manuela R. Zimmermann, Victor Lo, Maximo Sanz-Hernandez, Frank Jiang, Ann H. Kwan, Eryn L. Werry, Tuomas P. J. Knowles, Michael Kassiou, Georg Meisl, Matthew H. Todd, Peter J. Rutledge, Margaret Sunde

## Abstract

Alzheimer’s disease is imposing a growing social and economic burden worldwide and effective therapies are required. Strategies aimed at the removal of fibrillar plaques formed by the amyloid-β peptide have not proved therapeutic and the focus has shifted to approaches that target the cytotoxic oligomeric amyloid-β species that are populated before fibrils are deposited. We have designed and synthesized perphenazine-cyclam conjugates that specifically and rapidly bind to the monomeric form of Aβ42, reducing the production of both cytotoxic oligomers and amyloid fibrils. We have applied detailed kinetic analysis and NMR spectroscopy to show that the perphenazine-cyclam conjugates divert the Aβ42 monomer into amorphous aggregates that are not toxic to differentiated SH-SY5Y cells *in vitro*. Unlike most other amyloid inhibitors studied to date, these conjugates inhibit oligomer and fibril assembly even in the presence of pre-formed fibrillar seeds, demonstrating that they act through a monomer sequestration mechanism. These modular, three-dimensional conjugates therefore effectively prevent monomer-dependent secondary nucleation, the autocatalytic process that generates the majority of toxic oligomers.

## Introduction

Alzheimer’s disease (AD) is the most common form of dementia and one of the most devastating diseases of the current age. There are currently no approved disease-modifying therapies for this neurodegenerative disease. Although the full sequence of critical molecular events that lead to neuronal death is yet to be defined, self-assembly of the amyloid-β peptide (Aβ) into soluble oligomers and amyloid fibrils has been shown to generate neurotoxic oligomeric species^1-3^. These oligomers impact neuronal activity through effects on cell membranes, calcium homeostasis, and glutamate reuptake, amongst other deleterious mechanisms^4-10^. The failure of clinical trials aimed at reducing the accumulation of mature Aβ fibrils in extracellular plaques has seen efforts shift towards intervention at earlier stages in the pathway, with the aim of inhibiting formation of these toxic, pre-fibrillar oligomers^11^.

Aβ amyloid formation is a complex, multiple-step process, initiated by primary nucleation from Aβ monomers. This is followed by recruitment of additional monomers, leading to the formation of multiple different soluble oligomers with a range of sizes and toxicities^12^. As amyloid fibrils form, they offer sites for elongation at their ends, and for secondary nucleation on their lateral surfaces^13, 14^, generating further oligomeric species. Monomer-dependent secondary nucleation is a significant source of cytotoxic oligomeric Aβ42 species and is the dominant nucleation process even in cerebrospinal fluid and the presence of phospholipid membranes^15^.

Effective inhibition of Aβ-associated toxicity is likely to require a mechanism that is effective against all steps of the pathway^16^. Several small-molecule compounds have been shown to suppress primary nucleation or to accelerate fibril formation^17-19^ and have shown promise in animal models but are yet to show efficacy in clinical trials^20^. Some small molecules increase the population of neurotoxic Aβ oligomers^21^ and other proposed inhibitors are not effective in the presence of pre-formed fibrils^17, 22^. The molecular chaperone domain pro-SPC and certain antibody fragments are inhibitors of secondary nucleation^23, 24^, as are some small molecules^18, 25^. The efforts to identify effective modulators of the Aβ assembly process have been hampered by the use of poorly characterised preparations of the peptide and by testing against Aβ40, the less amyloidogenic and less clinically relevant isoform, rather than Aβ42 ^26-28^. The growing impact of AD necessitates continued efforts directed at the development of therapeutic modulators.

We have shown previously that cyclam derivatives with triazole-linked pendant groups can adopt a range of different, defined pendant geometries depending on the connectivity of the triazole linker and the metal ion coordinated by the macrocycle^29-31^. Combining these insights with the documented capacity of polymeric perphenazine conjugates to modulate Aβ aggregation^28^, we designed a series of molecules in which cyclam is linked via click-derived triazoles to pendant groups that are known to interact with Aβ42.

Quantitative kinetic analysis demonstrates that this series of molecules effectively and rapidly sequesters the monomeric form of Aβ42. We show that this monomer sequestration activity inhibits all downstream events that are dependent on monomer concentration, including suppression of the formation of toxic oligomers and amyloid fibril assembly, even in the presence of preformed fibril seeds. These novel molecules redirect the Aβ42 formation pathway towards the generation of amorphous aggregates that are not toxic to differentiated, neuron-like SH-SY5Y cells. This study provides the first demonstration of a reduction in neurotoxicity resulting from the designed diversion of the Aβ42 monomer into non-toxic oligomers.

## Results

### Perphenazine-cyclam conjugates inhibit Aβ42 amyloid assembly

A series of cyclam conjugates was designed to enable a systematic investigation of the Aβ aggregation and amyloid assembly pathways. Six parent cyclam derivatives were prepared: the mono-substituted naphthyl (**C1**), dopamine (**C4**) and perphenazine (**C7**) analogues, and corresponding bis-substituted compounds with naphthyl (**C10**), dopa (**C13**) and perphenazine (**C16**) pendants, along with their zinc(II) and copper(II) complexes (mono-naphthyl **C2, C3**; mono-dopa **C5, C6**; mono-perphenazine **C8, C9**; bis-naphthyl **C11, C12**; bis-dopa **C14, C15**; and bis-perphenazine **C17, C18**) (Fig. 1A and SI Fig. S1A). Initial screening of these compounds showed the bis-perphenazine derivative (ligand **C16)** to have promising capacity to suppress amyloid formation by both Aβ40 and Aβ42 (SI, Fig. S1B and C).

**Fig. 1.**
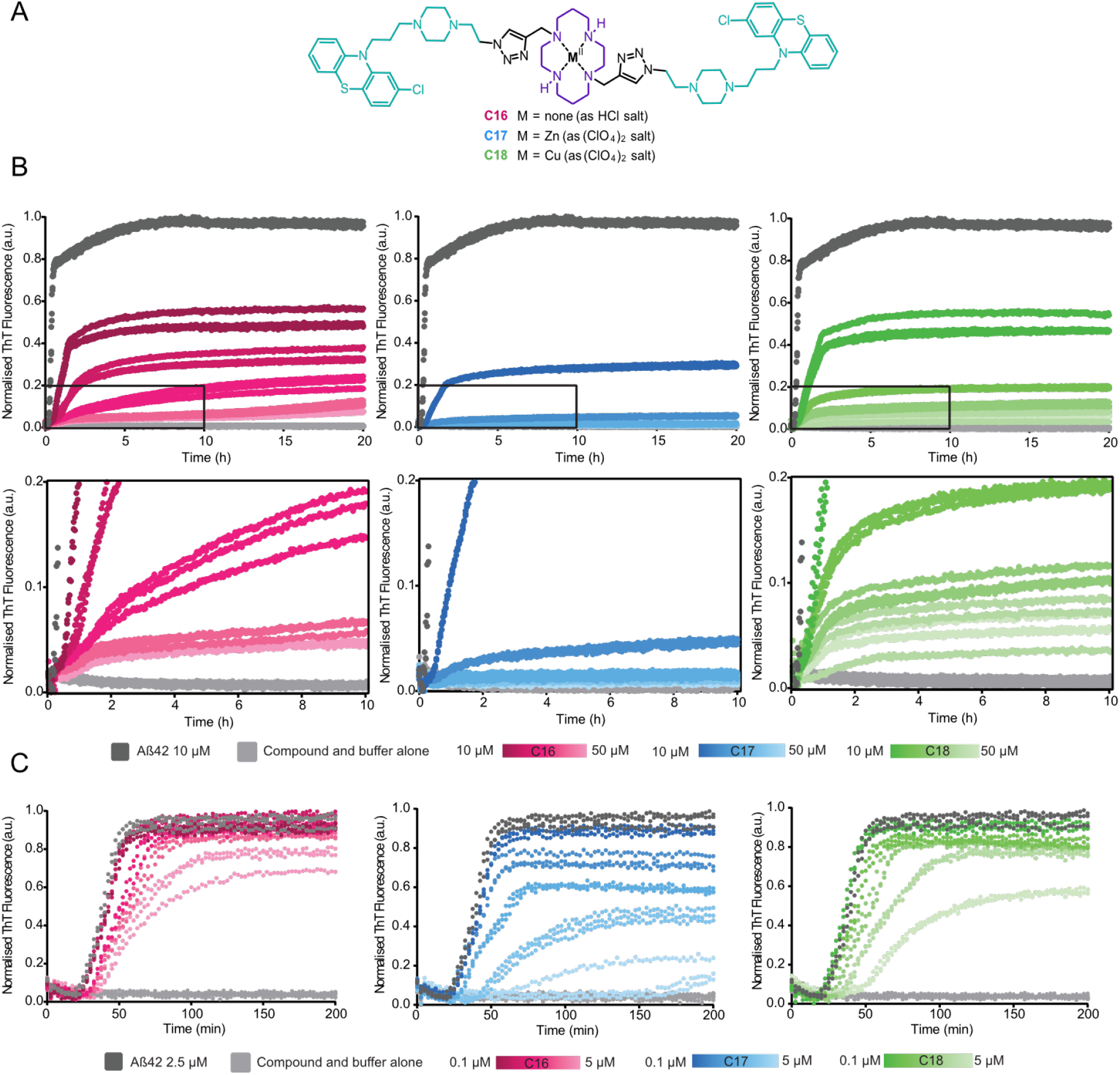
Time courses of fibril formation by Aβ42 show concentration-dependent inhibition of aggregation by **C16, C17** and **C18**. *(A)* The lead compounds used in this study combine a cyclam macrocycle (purple) with pendant perphenazine moieties (cyan), conjugated via a click-derived triazole (black), and deployed either as the unmetallated HCl salt **C16**, the zinc perchlorate complex **C17**, or the copper(II) perchlorate **C18**. *(B)* 10 μM monomeric Aβ42 in the absence or presence of either 1% DMSO (dark grey) or 10 – 50 μM of **C16** (pink), **C17** (blue) and **C18** (green). Lower section displays an enlarged 10-hour window. N = 3 (SI Fig S2). *(C)* 2.5 μM monomeric Aβ42 in the presence of either 1% DMSO (dark grey) or 0.1, 1, 2, 3 or 5 μM of compounds. N = 4 (SI Fig S3). All samples were measured in triplicate under quiescent conditions at 37 °C. Traces have been corrected for time zero and data normalised as a function of the maximum fluorescence intensity, using GraphPad Prism.

We prepared the bis-perphenazine ligand (**C16**) and its zinc (**C17**) and copper (**C18**) analogues and determined the ability of these three cyclam conjugates to inhibit amyloid formation by Aβ42 (SI, Fig. S1C,D). We have characterised the inhibitory activity of the molecules against Aβ42 amyloid assembly. This isoform is clinically associated with an increase in amyloid deposition^32^ and we have utilised highly purified, monomeric recombinant Aβ42, prepared using the methods of Linse and colleagues^33, 34^. Aβ42 prepared according to these protocols displays highly reproducible kinetics of amyloid assembly. The three cyclam conjugates **C16, C17** and **C18** were found to be potent inhibitors of Aβ42 amyloid formation, as monitored by ThT fluorescence (Fig. 1; SI Fig. S2 and Fig. S3).

The inhibitors demonstrated a concentration-dependent effect, and at a 1:5 molar ratio of Aβ42:compound, using 10 µM Aβ42, amyloid assembly was almost completely suppressed, as judged by the reduction in the level of Thioflavin T (ThT) fluorescence observed (Fig. 1B). Assays at lower Aβ42 concentration, using a range of inhibitor concentrations that allowed the plateau phase of amyloid assembly to be reached, revealed that all three compounds also introduce an increase in lag phase. The length of the lag phase increases with increasing inhibitor concentration (Fig. 1C). The zinc(II) complex **C17** displayed the strongest inhibition, followed by copper(II) complex **C18**, and the uncomplexed ligand **C16**, highlighting the capacity of the central metal ion to influence the three-dimensional shape of the inhibitor – in particular the relative orientation of the two pendant groups ^29, 30, 35^ – and resulting impact on the mechanism of Aβ42 amyloid inhibition. The exact nature of this inferred variation in shape between **C16, C17** and **C18** remains to be elucidated.

Perphenazine alone does not display strong inhibition of Aβ42 amyloid formation under these conditions, highlighting the fact that it is not the perphenazine moiety *per se* that is responsible for the inhibitory activity (SI Fig. S4). The observed inhibition is also not due to an effect of unchelated metal ions directly on Aβ42. Previous studies have demonstrated that Zn(II) and Cu(II) cations can themselves inhibit Aβ42 amyloid formation^36-38^, reflected by an increase in the lag phase or a decrease in the observed ThT intensity (SI Fig. 5A). However, we observed that **C17** and **C18** remain effective inhibitors of Aβ amyloid formation even in the presence of the chelating agent EDTA, which would sequester any free metal ions and block their inhibitory capacity (SI Fig. 5B).

### The perphenazine conjugates divert Aβ42 from amyloid assembly towards amorphous aggregates

A sensitive protein concentration assay confirmed that when the plateau of assembly was reached, in samples containing Aβ42 only, or Aβ42 in the presence of a 5-fold molar excess of inhibitor, all Aβ42 was converted to an insoluble form that was readily pelleted by centrifugation (SI Fig. S6). Transmission electron microscopy was used to examine the final morphology of the Aβ42 material present after ∼40 h incubation with compounds or 1% DMSO as control. An inhibitor concentration dependent shift from the characteristic fibrillar morphology of Aβ42 amyloid (Fig. 2A) to amorphous aggregates (Fig. 2B, C & D) was observed. The amorphous aggregates had a beads-on-a-string appearance, with the “beads” approximately 20 nm wide and the “strings” >1 µm in length. Only amorphous material was observed when Aβ42 was incubated in the presence of **C16**, even at 1:1 molar ratio of **C16**:Aβ42, while a combination of fibrillar and amorphous material was observed in samples prepared at 1:1 molar ratios of **C17**:Aβ42 and **C18**:Aβ42 (SI Fig. S7).

**Fig. 2.**
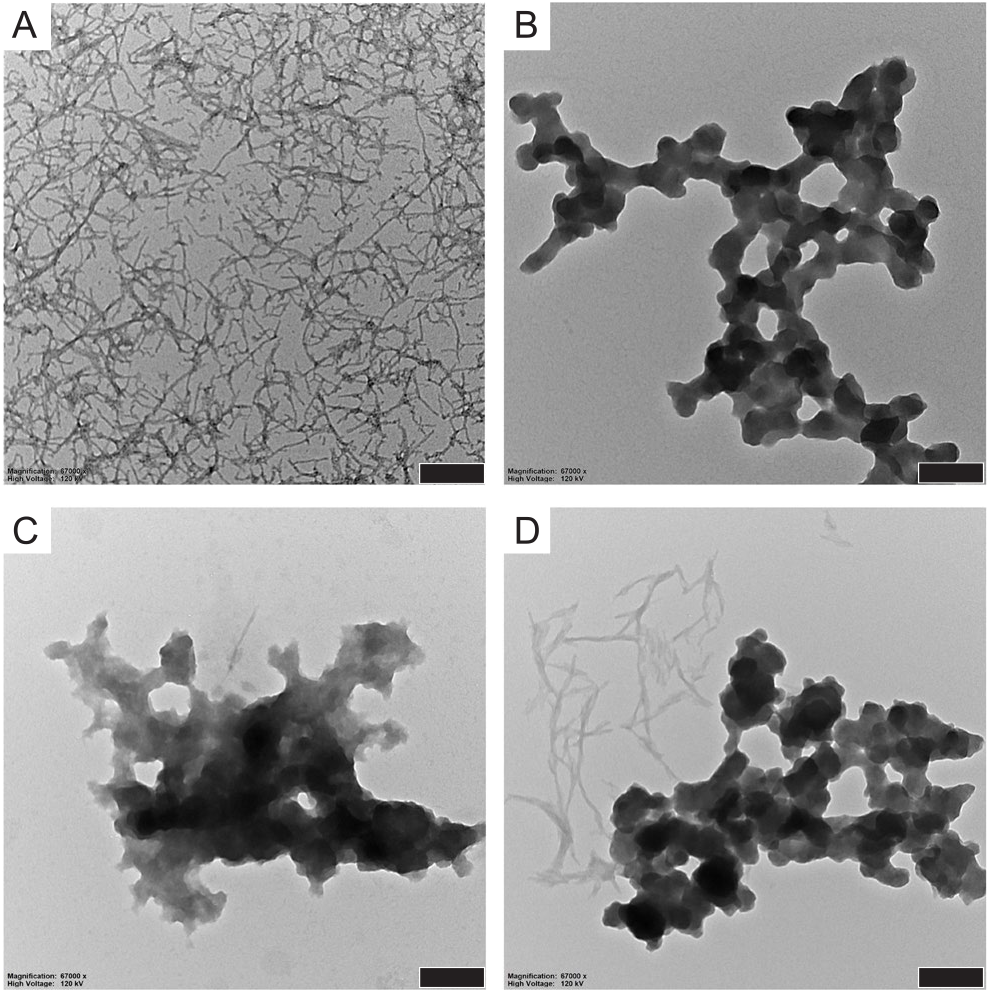
Transmission electron microscopy images showing representative examples of the morphology of fibrils formed by 10 μM Aβ42 alone (A) or aggregates formed by 10 μM Aβ42 in the presence of 50 μM **C16** (B), **C17** (C) and **C18** (D). Scale bar represents 200 nm.

### The perphenazine conjugates reduce neurotoxicity associated with Aβ oligomers

Many groups have demonstrated that soluble intermediate species formed by Aβ42 *in vitro* are neurotoxic^1-3, 39^. Accordingly, the ability of compounds **C16**–**C18** to prevent the formation of toxic Aβ42 species was indexed using SH-SY5Y cells which had been differentiated to an AD-appropriate phenotype by treatment with retinoic acid and brain-derived neurotrophic factor^40^. Lyophilised, monomeric Aβ42 was treated with hexafluoro-2-propanol (HFIP), lyophilised, and then resuspended in PBS, before being diluted in cell media and added immediately to cells. Aβ42 fibrils were prepared by incubating monomeric Aβ42 solution in cell media at 37 °C for 24 hours, a time point significantly past the plateau level in unshaken ThT experiments using cell media (SI Fig. S8). The compounds were added to differentiated SH-SY5Y cells alone or added together with Aβ42 in a monomeric or fibrillar form. After 24 hours, the number of metabolically-active cells was measured using a CTB fluorescence assay. The CellTiter-Blue^®^ (CTB) assay of cell viability was chosen, given published concerns regarding the use of MTT with Aβ42^1^.

By themselves, compounds **C16, C17** and **C18** displayed no significant toxicity towards differentiated SH-SY5Y cells, relative to vehicle only (Fig. 3A). Exposure of these cells to freshly prepared monomeric Aβ42 resulted in significant (P < 0.001) toxicity compared to vehicle, as expected due to the generation of oligomeric species, while the addition of **C16, C17** or **C18** led to significant protection against this neurotoxicity (Fig. 3B). Perphenazine was not cytotoxic by itself and imparted a similar level of protection as **C16, C17** and **C18**. These results contrast with the observed neurotoxicity of epigallocatechin gallate (EGCG) at the same concentrations and the lack of protection by this polyphenol against cytotoxicity induced by addition of monomeric Aβ42 (Fig. 3A,B). When Aβ42 was prepared in fibrillar form and then added to cells, no cytotoxicity was observed with fibrils alone or in the presence of compounds (Fig. 3C). Following from these experiments, independent concentration response assays were performed with **C16**. The IC_50_ for **C16** was determined to be 2.33 μM (n = 3; SD = 0.89) (Fig. 3D). Here we have used an established assay of neurotoxicity, with differentiated SH-SY5Y cells and well-characterised preparations of monomeric and fibrillar Aβ42. Although the origins of the neurotoxicity resulting from the conversion of Aβ42 from monomeric to fibrillar form *in vitro* are not defined, these results provide clear evidence that the generation of amorphous Aβ42 aggregates in the presence of **C16**–**C18** prevents the formation of neurotoxic species.

**Fig. 3.**
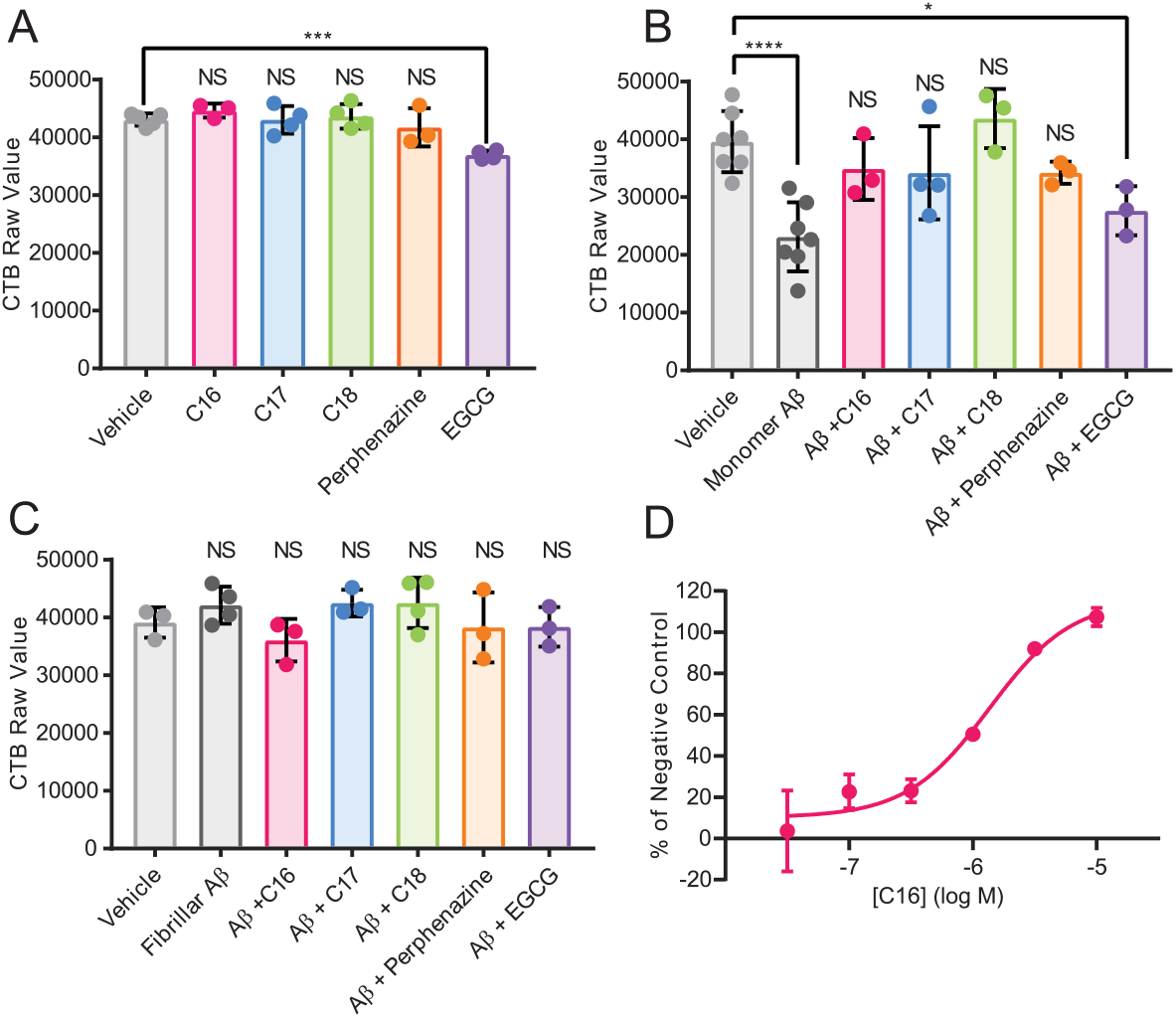
Cell viability was measured using the CellTiter Blue® assay after 24 h incubation with vehicle or 10 μM compound, with or without 20 μM Aβ42. (*A)* Compounds alone. *(B)* Compounds incubated with monomeric Aβ42. *(C)* Compounds incubated with fibrillar Aβ42. The figure displays the mean ± SD of N ≥ 3 independent experiments. One-way ANOVA followed by a Bonferroni’s multiple comparisons test at the 0.05 level was used to determine the difference between each condition and vehicle control (**** p < 0.0001, *** p < 0.001, * p < 0.05, NS = non-significant). *(D)* Representative concentration-response curve for **C16** incubated with Aβ42 showing mean ± SD of duplicates. IC_50_ = 2.33 μM (SD = 0.89). N = 3.

### C16, C17 and C18 inhibit Aβ amyloid formation through sequestration of Aβ42 monomers

The microscopic processes underlying amyloid formation of Aβ42 are primary nucleation, elongation and secondary nucleation^41, 42^. Their relative importance can be quantified through their rate constants, obtained by fitting integrated rate laws within the framework of chemical kinetics^43^. Data measured at 2.5 µM and 10 µM Aβ42, in the presence of a range of concentrations of inhibitor, were analysed using the online fitting software AmyloFit. The rate constants of the unperturbed systems agreed well with similar experiments^13^. To probe the mechanism of action of the inhibitory compounds **C16, C17** and **C18**, we varied systematically one rate constant in the rate law, or the level of available monomer, respectively^43^. By comparing how well these modified rate laws can describe the data, a mechanism that primarily involves inhibition of primary nucleation could be ruled out for **C16, C17** and **C18** (Fig. 4A, B and SI Fig. S9-13) as the origin of the changes in the kinetic behaviour in the presence of the inhibitor.

**Fig. 4.**
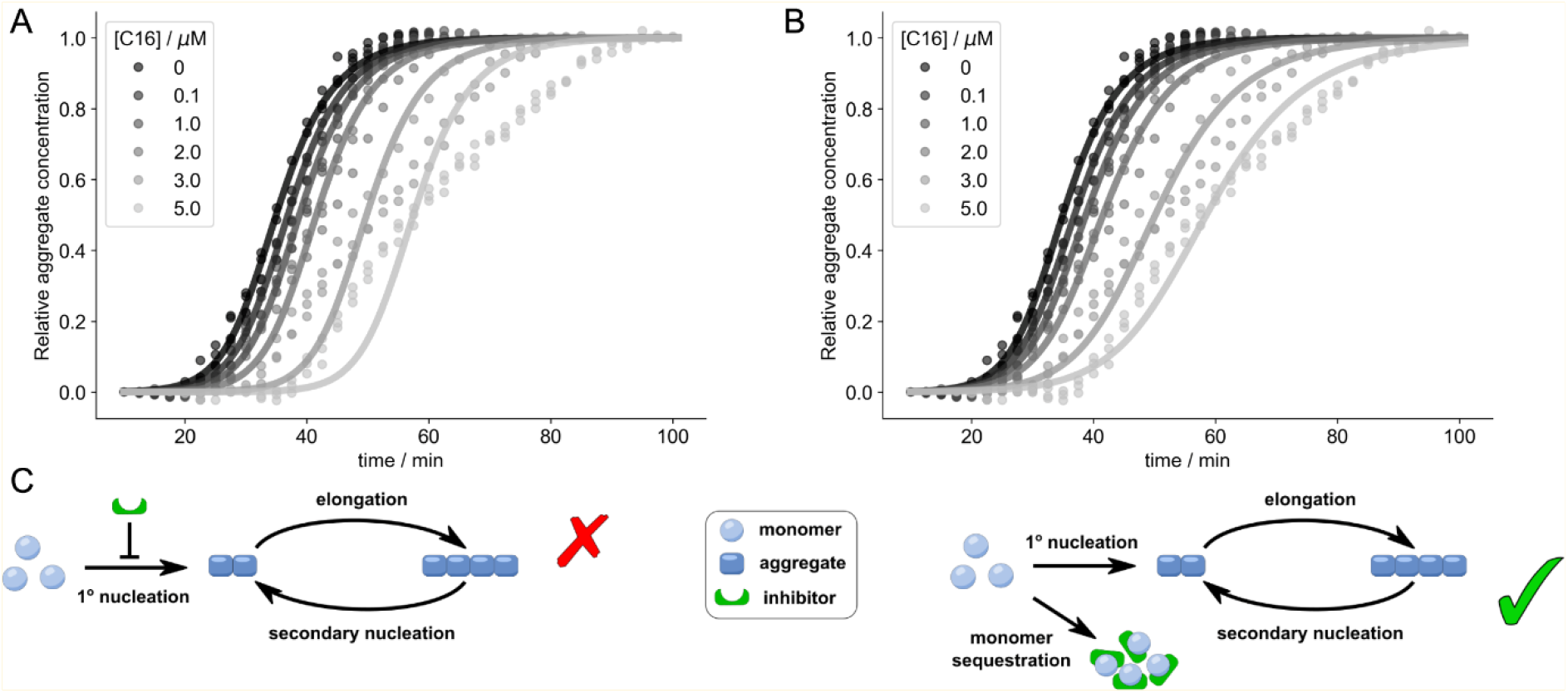
Chemical kinetics analysis reveals inhibition through monomer sequestration. To elucidate the major mechanism of inhibition, normalised aggregation curves measured in thioflavin T fluorescence assays were fitted under the constraint that deviation from the unperturbed system was allowed in only one microscopic step or the free monomer concentration, respectively. Representative curves shown in A and B, all fits shown in the SI. *(A)* The observed inhibition cannot be described through inhibition of primary nucleation. *(B)* Inhibition through monomer sequestration is a plausible mechanism based on the kinetic profiles. *(C)* Microscopic steps underlying amyloid formation, with the suggested mode of inhibition through monomer sequestration.

The introduction of small numbers of preformed fibril seeds at the beginning of the aggregation reaction can be used to bypass primary nucleation. These compounds remain effective as inhibitors of Aβ42 amyloid fibril formation even when exogenous fibril seeds are present (Fig. 5). In the presence of 2% or 5% seeds, and the presence of **C16–C18** at 1:2.5 or 1:5 Aβ42:compound molar ratio resulted in inhibition of Aβ42 assembly, similar to the level of inhibition observed in the absence of pre-formed seeds (Fig. 5 and SI Fig. S14). In addition to ruling out the blocking of primary nucleation as the source of the inhibitory effect, the results observed in the presence of seeds, when the aggregation reaction reaches completion much sooner, suggest that binding of the compounds to the monomeric form of Aβ42 occurs rapidly.

**Fig. 5.**
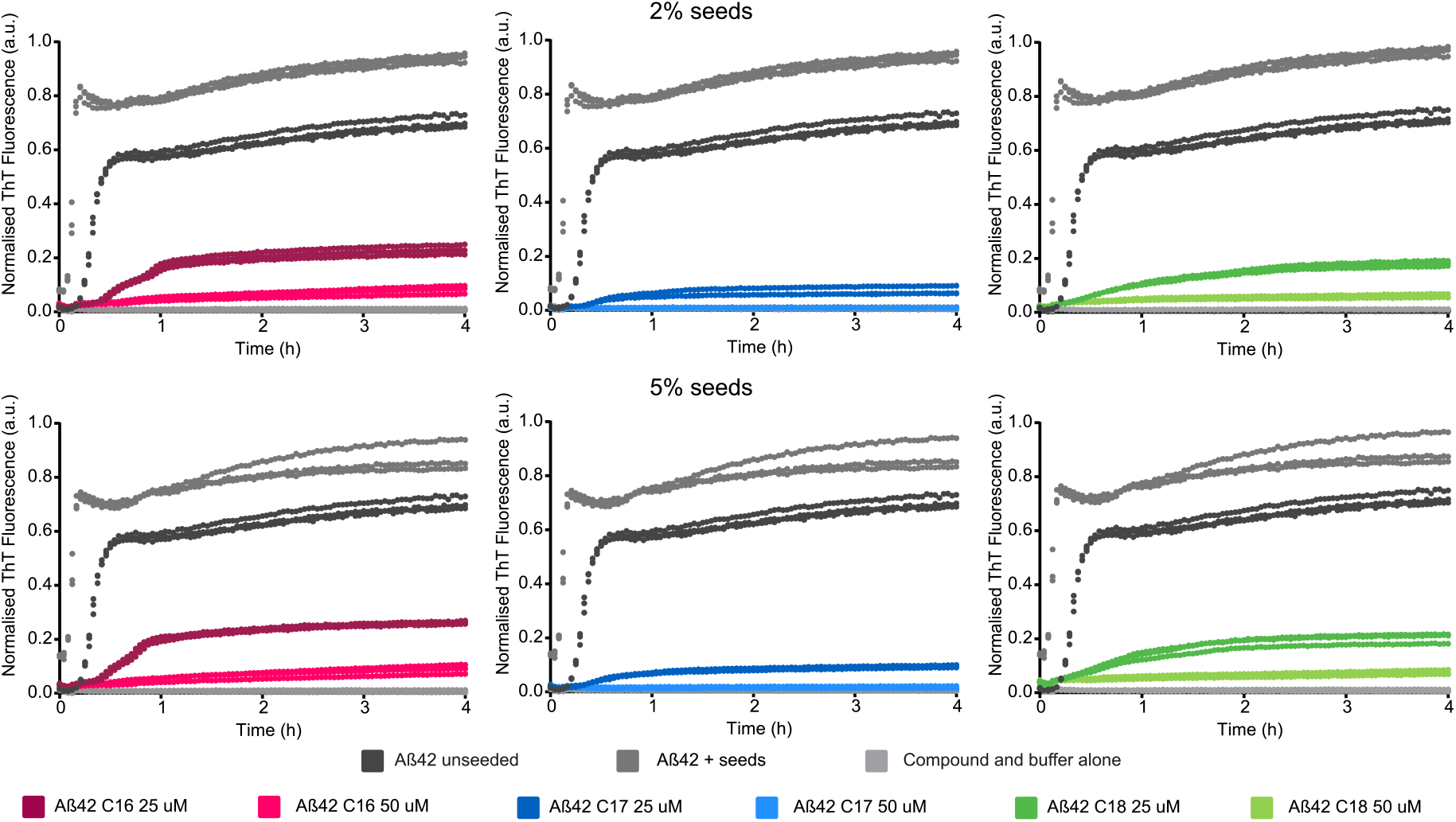
Characterisation of the effects of compounds on seeded Aβ42 aggregation. 10 μM monomeric Aβ42, in the absence of seeds (black) or presence of 2% (top panel) or 5% (bottom panel) seeds and 1% DMSO (dark grey), 25 or 50 μM of compounds. N = 3 (SI Figs S14). All samples were measured in triplicate under quiescent conditions at 37 °C. Traces have been corrected for time zero and data normalised as a function of the maximum fluorescence intensity, using GraphPad Prism.

Based on the fits to the normalised kinetic data alone, mechanisms based on inhibition of secondary nucleation, inhibition of elongation, and monomer sequestration were similarly likely as the main modes of action of compounds **C16**–**C18** (SI Fig. S9-13). However, we found a clear correlation between the ThT intensity at completion of the aggregation reaction and the effective monomer concentration determined from the fits (Fig. 6A, B, C insets). As the fitting was performed on normalised kinetic data, these two quantities constitute independent measures of the same physical property. The correlation between an equilibrium measure, the plateau level, and a parameter estimated on the basis of normalised kinetics, the effective monomer concentration, provides further independent evidence for monomer sequestration by these inhibitors. In addition to accounting for the delay in aggregation and decrease in ThT signal, monomer sequestration by the inhibitors also provides a mechanistic explanation for the appearance of amorphous material as discussed above. These fits were made assuming that this monomer sequestration proceeds much faster than aggregation and can thus be modelled by a reduced effective monomer concentration.

**Fig. 6.**
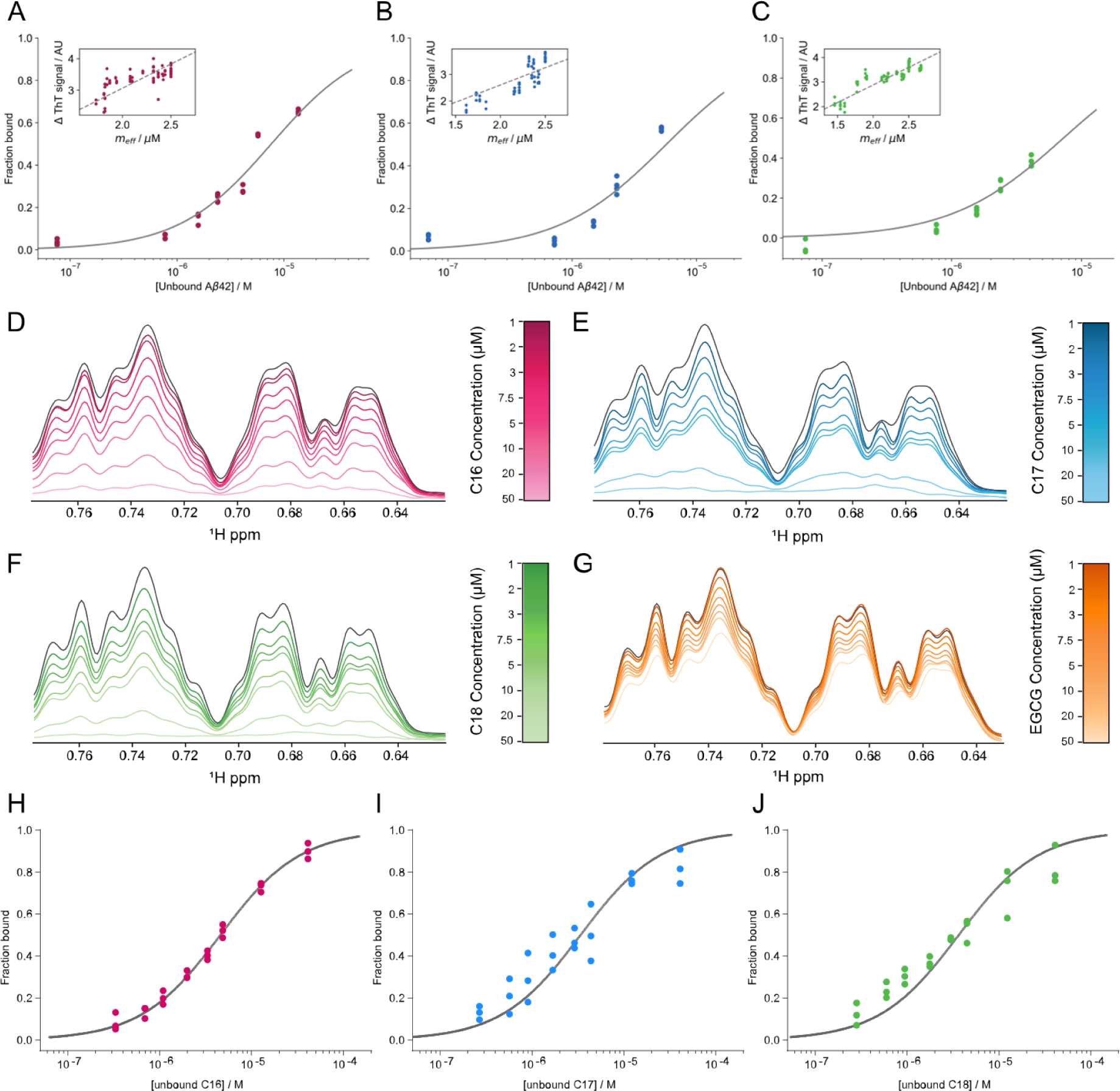
Dissociation constants determined from ThT assays and ^1^H NMR. Monomer-inhibitor binding curves for *(A)* **C16**, *(B)* **C17** and *(C)* **C18**, with correlation between the effective free monomer concentration and the absolute increase in ThT signal shown as insets. One-dimensional ^1^H NMR spectroscopy to determine apparent *K*_*D*_ for compounds. Unlabelled 10 μM Aβ42 (dark grey) was incubated with 1–50 µM *(D)* **C16**, *(E)* **C17**, *(F)* **C18** and *(G)* **EGCG**, and spectra recorded at 4 °C. Methyl proton region is displayed, all regions analysed are shown in SI Fig. 15. Increasing concentration of **C16**–**C18** resulted in a decay in protein signal intensity. Binding curves for *(H)* **C16**, *(I)* **C17** and *(J)* **C18** constructed from signal decay curves and used to calculate the apparent *K*_*D*_.

Further support for monomer sequestration as the dominant mode of inhibition of Aβ42 aggregation by **C16, C17** and **C18** was gained by obtaining consistent values for the apparent dissociation constant of monomer and inhibitor, *K*_*D*_, by orthogonal methods. First, the effective monomer concentration obtained through the chemical kinetics analysis at different monomer and inhibitor concentrations was used to construct monomer-inhibitor binding curves (Fig. 6A, B, C). To fit these data, we assumed a simple one-to-one binding, yielding *K*_*D*_ values of 7.6 μM (95% CI, 6.6–9.1) for **C16**, 5.7 μM (95% CI, 4.6–7.1) for **C17** and 7.4 μM (95% CI, 6.2–9.1) for **C18**.

Secondly, we utilised one-dimensional ^1^H NMR spectroscopy to determine an apparent binding constant for the interaction between the inhibitors and Aβ42. For each of **C16, C17** and **C18**, inhibitor was titrated into a solution containing 10 μM unlabelled Aβ42, over the inhibitor concentration range of 1–50 μM. A decay profile in the protein signal intensity across the spectrum was observed with increasing concentration of all three inhibitors (Fig. 6 and SI Fig. S15). A 5:1 ratio of inhibitor:Aβ42 resulted in a reduction of proton signals to baseline, consistent with the sequestration of monomer into very large and/or insoluble assemblies, the precipitation of material during the course of the NMR titrations, and the amorphous aggregates observed by TEM (Fig. 3). The signal decay curves were fitted to a quadratic equation for the calculation of the apparent *K*_*D*_ of each compound, yielding 4.6 µM (95% CI, 3.1–6.5) for **C16**, 3.4 µM (95% CI, 2.2–4.9) for **C17** and 3.7 µM (95% CI, 2.4–5.3) for **C18**. The observed effect of these perphenazine cyclam conjugates on Aβ42 assembly is different to that of the polyphenol EGCG, where the presence of EGCG results in only a small decrease in the intensity of the observed 1H spectrum from Aβ42, up to 31% signal loss at a 5-fold excess of small molecule (Fig. 6D and SI Fig. S15). This is evidence that upon addition of EGCG, the monomeric Aβ42 species is still the most prevalent in the sample. In contrast, when the less aggregation-prone Aβ40 interacts with EGCG, specific interactions are observed by solution NMR and it forms insoluble oligomers that are in equilibrium with soluble Aβ40^44^.

A monomeric sample of recombinant ^15^N-labeled Aβ42 was prepared and analysed in the absence and presence of **C16** to probe the regions of the polypeptide involved in the interaction. The ^15^N HSQC spectrum collected in the absence of any added compound reflected the high quality of the recombinant, monomeric Aβ42 sample and the chemical shifts were superimposable upon those reported by others previously for Aβ42^45^ (SI Fig. S16). Addition of **C16** resulted in an immediate reduction in signal intensity that was uniform across all peaks in the spectrum. Protein that is not sequestered into an amorphous aggregate remains in the solution at the 1:1 ratio of protein:inhibitor. No peak-specific changes in signal intensity or chemical shift were detected that could be attributed to binding of **C16** to a distinct region of monomeric Aβ42. Instead, the loss of signal is consistent with the binding of monomer to **C16** resulting in the generation of a conformation that is highly aggregation-prone and rapidly forms large structures, undetectable by solution NMR.

## Discussion

We have designed a class of compounds that specifically sequester the monomeric form of Aβ42 into non-fibrillar assemblies that are different to the cytotoxic oligomeric forms associated with Aβ42 amyloid formation. The process by which monomeric Aβ42 converts to the mature fibrillar form detected in amyloid plaques involves multiple oligomeric and prefibrillar species that may act as reactants, products, intermediates, and catalysts of the process and contribute to toxicity to different extents. The ability to interrogate a change in aggregation kinetics caused by an inhibitor, and thereby identify the mechanism of inhibition, offers new opportunities to understand the consequences of intervention at defined points within the Aβ42 assembly pathway. Many studies of peptide-based inhibitors that incorporate “disruption elements” report success in preventing the formation of the β-structure that stabilises amyloid fibrils and can influence elongation^46^ and inhibitors have been described that specifically inhibit secondary nucleation^18^, but the impact of these interventions on residual Aβ monomer concentration has not always been described. The kinetic theory of protein aggregation inhibition reveals that inhibitors such as the cyclam-perphenazine conjugates described here, which sequester monomeric Aβ42, will reduce the rates of primary nucleation, elongation and secondary nucleation, and are therefore likely to have a large impact on amyloid assembly overall^16^.

In addressing AD the sequestration of monomer into non-toxic assemblies and consequent reduction of oligomer formation, could be beneficial. Studies of the effect of EGCG on Aβ42 indicate that it converts the peptide to off-pathway oligomers and remodels pre-formed fibrils, although these effects are not rapid^26, 47, 48^. The intrinsically disordered and dynamic nature of monomeric Aβ42 has impeded the use of traditional approaches that focus on identifying chemical moieties that bind to defined sites on the target. Recently a small molecule (10074-G5) that binds to the monomeric form of Aβ42 has been reported^16, 22^.The binding of 10074-G5 to Aβ42, with *K*_*D*_ of ∼40 µM, sequesters Aβ42 in its soluble form and reduces its hydrophobicity. In contrast, we observed an immediate uniform reduction in signal intensity across the spectrum when **C16** was added to Aβ42 at a 1:1 ratio, indicating that **C16** sequesters Aβ42 in a polymeric, insoluble assembly. The small molecule 10074-G5 reduces functional deficits in a *C. elegans* model of Aβ42-associated toxicity, however its impact on neuronal toxicity is yet to be reported and it does not display a concentration-dependent inhibition of seeded assembly *in vitro*. Thus, further work is required to define the relative benefit of Aβ42 sequestration in monomeric versus non-toxic oligomeric form.

The inventive modular design of the compounds described here offers a toolkit for future elucidation of the key moieties and properties that influence Aβ42 monomer binding and subsequent formation of amorphous aggregates. Compounds that were effective at Aβ42 monomer sequestration protected against Aβ42-induced cytotoxicity. This scaffold provides a base for future rational design of more potent compounds that act through the same mechanism, yet which are accessible via a divergent synthetic strategy. This class of compounds can provide wide mass-to-charge and hydrophobic-to-hydrophilic surface area ratios, while the attachment of both rigid and flexible pendants provides a great diversity of possible interactions with client protein targets. The switchable geometry allows maintenance of a constant periphery while varying the gross molecular shape. Given that self-assembly and aggregation of proteins are three-dimensional processes, this level of structural control has significant potential to contribute to effective inhibition. AD is one of over 50 human disorders considered to be proteinopathies. The critical pathogenic species associated with protein aggregation and amyloid formation in each of these disorders likely have unique features. However, a similar dissection of assembly pathways with modular scaffolds decorated with protein-specific binding moieties may reveal points for therapeutic intervention in other protein aggregation-related diseases.

## Materials and Methods

### Synthesis of lead compounds C16–C18

Full synthetic procedures and characterisation data are detailed in the Supporting Information.

### Reaction scheme 1

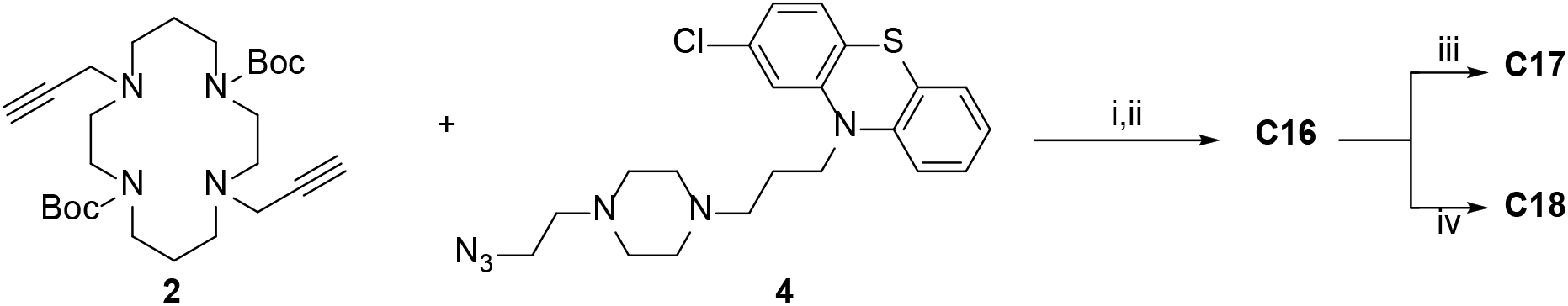

Cyclam (1,4,8,11-tetraazacyclotetradecane) **1** was converted to the di-Boc-di-propargyl derivative **2** in 68% yield over four steps using previously reported methods^49, 50^. Perphenazine **3** was converted to the azide **4** (79%) by activating its primary alcohol with diphenylphosphoryl azide (DPPA) and displacing with sodium azide in dimethylformamide (DMF). The azide **4** (2 equivalents) and bis-alkyne **2** were coupled using a copper-catalysed azide/alkyne cycloaddition reaction (CuAAC) to yield the protected ligand **5** (46%), before the Boc groups were removed using 4 M HCl in dioxane to afford **C16** (74%). The HCl salt **C16** was converted to its zinc(II) (**C17**) and copper(II) (**C18**) complexes by first treating with Ambersep 900 hydroxide form resin to generate the free amine, then stirring overnight at room temperature with either zinc(II) perchlorate hexahydrate (for **C17**, 63%) or copper(II) perchlorate hexahydrate (for **C17**, 64%). All compounds were dissolved in 100% DMSO and stored at 4 °C until use. Reagents and conditions: i. CuSO_4_.5H_2_O (0.2 equiv.), sodium ascorbate (0.5 equiv.), THF/H_2_O (2:3), rt, 16 h, 46%; ii. 4 M HCl in dioxane (excess), 0 °C, 1 h, 74%; iii. Ambersep 900 hydroxide form resin, MeOH, 15 min, then Zn(ClO_4_)_2_, MeOH, reflux, 16 h, 63%; iv. Ambersep 900 hydroxide form resin, MeOH, 15 min, then Cu(ClO_4_)_2_, MeOH, reflux, 16 h, 64%.

### Aβ42 Overexpression and purification

Overexpression and purification of unlabelled Aβ42 was performed according to Walsh and colleagues^34^. Briefly, large cultures of *Escherichia coli* BL21 Gold (DE3) were incubated at 37 °C with shaking, induced with 0.5 mM IPTG and harvested by centrifugation. Purification involved a series of sonication and centrifugation steps followed by resuspension of inclusion bodies in 8M urea. Anion exchange chromatography using diethylaminoethyl cellulose beads was performed and protein eluted with increasing concentrations of NaCl (up to 150 mM). Aβ42 elution was assessed by electrophoresis, using by Novex™ 10–20% Tricine SDS-PAGE gels. Fractions containing elevated levels of Aβ42 were combined and filtered through a 30 kDa molecular mass cut-off centrifugal filtration device (Amicon^®^ Ultra-15). The combined sample was concentrated using a 3 kDa molecular mass cut-off centrifugal filtration device (Amicon^®^ Ultra-15), snap frozen, lyophilised and stored at –20 °C. ^15^N Aβ42 expression was performed according to Habchi and colleagues^18^, with purification as above.

### Size exclusion chromatography

An aliquot of lyophilised Aβ42 protein was solubilised in 6 M GuHCl until no large aggregates were visible. The sample was centrifuged at ∼3300 x *g* for 3 min, bath sonicated for 5 min in ice-water and centrifuged again at 16000 x *g* for 10 min at 4 °C. 500 μL of sample was drawn up carefully with a syringe and loaded onto a Superdex™ 75 Increase 10/300 GL size exclusion column running on an ÄKTA FPLC system. Protein was eluted at 0.5 mL/min in 50 mM ammonium acetate pH 8.5. Fractions (0.5 mL) were collected in Protein LoBind Eppendorf tubes and immediately placed on ice. Fractions were analysed for Aβ42 content using SDS-PAGE. Concentration was measured via spectrophotometer (ε275 = 1450). Fractions containing Aβ42 were combined and either used immediately or snap frozen and lyophilised.

### Thioflavin T assays

Protein samples (Aβ42) were prepared in triplicate in non-binding 96-well half area plates (Corning 3881) in 25 mM or 50 mM NaH_2_PO_4_ pH 7.4, 40 μM ThT to a final volume of 80 μL. Plate preparation was performed at 4 °C to minimise aggregation. Samples were incubated at 37 °C in a POLARstar Omega microplate reader (BMG Labtech), with excitation at 440 nm and the fluorescence emission recorded at 480 nm under quiescent conditions.

### Seeding

Seeds were prepared by incubating 10 μM Aβ42 in 50 mM NaH_2_PO_4_, pH 7.4 at 37 °C for 5 h. Protein solution was subject to 6 × 10 s sonication with 20 s rest on ice between cycles. The seeds were maintained on ice and used immediately.

### Enhanced BCA assay details

Kinetic assays were run to completion as judged by the plateau in ThT fluorescence (∼5 h). Contents of wells were removed and centrifuged at 16000 x *g* for 10 min to pellet fibrils and aggregates. Protein concentration was determined using a Pierce^®^ Bicinchoninic Acid Protein Assay Kit according to manufacturer’s enhanced protocol. Monomeric Aβ42 and compounds alone were used as controls.

### TEM

Samples were prepared on carbon-coated copper grids (200 mesh) with formvar support film (ProSciTech Pty Ltd.) that had been exposed to UV light (230 nm) for 10 min before sample application and stained with 2% aqueous uranyl acetate solution. Samples were examined with a FEI Tecnai T12 electron microscope operating at 120 kV and images captured with a Veleta CCD camera and processed using RADIUS 2.0 imaging software (EMSIS GmbH).

### Preparation of Aβ42 for cell assay and NMR

Lyophilised, purified protein was resuspended in HFIP at 1 mg/mL and incubated for 30 min at room temperature. The protein was snap frozen, lyophilised, and stored at –80 °C until use.

### Cell Culture and Differentiation

SH-SY5Y cells were cultured in in Dulbecco’s Modified Eagle Medium: Nutrient Mixture F-12 (DMEM/F12; Invitrogen, Carlsbad, CA, USA) supplemented with 10% fetal bovine serum (FBS; Gibco, Uxbridge, UK) at 37 °C and 5% CO_2_. Cells were fed every 2-3 days and passaged using Trypsin-EDTA (0.25%). Cells were detached using Trypsin-EDTA, centrifuged (5 min, 125 x *g*), counted and then plated in a Cell-bind 96-well plate at a density of 0.5 × 10^4^ cells/well in 10 μM all-trans retinoic acid (RA; Sigma-Aldrich) in DMEM/F12 with 10% FBS. Media was replaced daily for 4 days (RA is light sensitive, so all steps involving RA were completed in the dark). After a 4-day incubation with RA, media was switched to DMEM/F12 (no serum) with 50 ng/mL BDNF (Sigma-Aldrich) for 3 days. After a total of 7 days of differentiation the cells were used for the experiments.

### Aβ42 preparation, compound treatment and CellTiter-Blue^®^ viability assay

24 h prior to the cells finishing differentiation, an aliquot of Aβ42 was centrifuged at 16,000 x *g* for 10 min and dissolved in PBS at 200 μM, then diluted to 20 μM in DMEM/F12 with 10% FBS. The solution was stored in the incubator (37 °C, 5% CO_2_) in a low-bind Eppendorf to promote Aβ42 fibrillization. After cell differentiation was complete, for the Aβ42 monomeric preparation, a second aliquot of Aβ42 was centrifuged at 16,000 x *g* for 10 min, dissolved in PBS at 200 μM and immediately diluted to a final concentration of 20 μM in DMEM/F12 (with 10% FBS). Experimental compounds (10 μM, 0.1%) or DMSO control were added to the fibrillar or monomeric Aβ42. Differentiation media was aspirated, 100 μL of each condition added to the cells and incubated for 24 h (37 °C, 5% CO_2_). In the monomeric preparation, the time between dissolving the Aβ42 and applying onto the cells was minimised (less than 2 min total). After 24 h, CellTiter-Blue^®^ (CTB; Promega G8080) (20 μL/well) was added and incubated for 4 h. The fluorescent emission at 590 nm was measured using a POLARstar Omega microplate-reader (BMG Labtech); viable cells metabolise this agent and reduce it to a fluorescent compound.

### Statistical Analysis

One-way ANOVA (analysis of variance) followed by a Bonferroni’s multiple comparisons test was used to determine the differences between mean CTB values of differentiated SH-SY5Y cells treated with Aβ42 and/or test compounds relative to vehicle control. Significance threshold was defined as p <0.05.

### Kinetic analysis

The kinetic data obtained in ThT assays were analysed using the online fitting software AmyloFit, following the cited protocol^43^. The model ‘secondary nucleation dominated, unseeded’ was used, with the reaction orders set to 2 as typical for Aβ42 and under the constraint of allowing deviations from the unperturbed kinetics in only one parameter at a time to investigate which microscopic step or which species of the aggregation mixture was most likely affected by the inhibitors. Repeats of the experiment were analysed separately and combined at the end to estimate uncertainty.

### *K*_*d*_ determination

To determine the value of the dissociation constant, *K*_*d*_, between free monomer and inhibitor, a one-to-one binding was assumed and the free monomer concentration, *m*, expressed as a function of the total inhibitor concentration, *I*_*tot*_, the total monomer concentration, *m*_*tot*_, and *K*_*d*_,

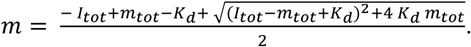

The free monomer concentration was estimated as the effective monomer concentration in the kinetic assays or the relative one-dimensional 1H NMR signal, respectively. Likelihoods were calculated using grid approximations, under the assumption of Gaussian noise.

### NMR / *K*_*d*_ Analysis

All NMR data were collected at 4 °C using unlabelled 10 μM Aβ42 or ^15^N-labelled Aβ42 in 20 mM NaH_2_PO_4_ pH 7.0, 10% D_2_O, 1 mM DSS. Individual titrations with **C16, C17, C18** or EGCG were carried out by incubating unlabelled 10 μM Aβ42 with increasing concentrations of the compounds, ranging from 0 to 50 μM. 9 titration points were recorded for each compound. The NMR signals from one-dimensional ^1^H spectra were used to monitor the binding. Each spectrum was recorded using 512 scans. A ^1^H–^15^N heteronuclear single quantum coherence (HSQC) spectrum was also recorded for ^15^N-labelled Aβ42 in the presence of 10 μM **C16**. The HSQC spectrum was recorded overnight, for 240 scans, with a spectral width of 25 ppm (^15^N offset at 117.5 ppm). All spectra were obtained using a 18.8 T (800 MHz ^1^H frequency) Advance III HD Bruker spectrometer equipped with a cryoprobe.

## Acknowledgements

The work was funded by Australian Research Council Discovery Project grant (DP200102463) to MS, AK and PJR. SRB, JSPA and MAS were supported by Research Training Program Scholarships from the Australian Government (Department of Education, Skills and Employment). MK, MAS and ELW’s research is supported by the National Health and Medical Research Council of Australia (NHMRC; APP1132524). MK is an NHMRC Principal Research Fellow (APP1154692). MRZ acknowledges support from the Herchel Smith Fund. The authors thank Elva (Meng) Shi and Dr Malcolm Spain for assistance with synthesis and characterisation of compounds and Dr Bill Bubb for comments on the manuscript. The authors acknowledge the facilities and the scientific and technical assistance of staff within the Sydney Analytical and Sydney Microscopy & Microanalysis Core Research Facilities at the University of Sydney. We acknowledge and pay respect to the Gadigal people of the Eora Nation, the traditional owners of the land on which we research, teach, and collaborate at the University of Sydney.

