## Supplementary data for "Perphenazine-macrocycle conjugates rapidly sequester the Aβ42 monomer and inhibit amyloid formation"

[1]School of Medical Sciences, The University of Sydney, Sydney, Australia. [2]School of Chemistry, The University of Sydney, Sydney, Australia. [3]Centre for Misfolding Diseases, Department of Chemistry, University of Cambridge, Cambridge UK. [4]Department of Life Sciences, Imperial College London, South Kensington, SW7 2AZ, UK. [5]School of Life and Environmental Sciences, The University of Sydney, Sydney, Australia. [6]Brain and Mind Centre, The University of Sydney. [7]Cavendish Laboratory, University of Cambridge, JJ Thomson Avenue, CB4 0HE, UK. [8]School of Pharmacy, University College London, London UK.

*Co-corresponding authors: Margaret Sunde, Peter Rutledge, and Matthew Todd

**This PDF file includes:**

Supplementary text
Figures S1 to S16
SI References

### Supplementary Methods

#### General Materials and Instrumentation

Anhydrous DMF was collected from a PureSolv MD7 solvent purification system with anhydrous alumina columns. All commercially available chemicals and solvents were purchased from Sigma-Aldrich, Merck Millipore, Fluka, Matrix Scientific or Combi-Blocks and used without further purification. Reactions were cooled to 0 °C with an ice bath. Automated flash column chromatography was performed on a Biotage Isolera Spektra One using Biotage SNAP KP-Sil cartridges (filled with Merck Silica Gel 60 (0.040-0.0630 mm)). Reversed phase chromatography was performed using either Biotage SNAP C18 or iLOK SL prepacked C18 columns. Analytical TLC was performed on Merck silica gel 60 F254pre-coated aluminium plates (0.2mm) and visualised with UV (254 nm and 280 nm) and subsequent staining with either potassium permanganate. $^{1}$H and $^{13}$CNMR spectra were obtained using either a Bruker Avance DPX200 (200 MHz and50 MHz respectively), DPX300 (300 MHz and 75 MHz) or DPX500 (500 MHz and 125 MHz) instrument and processed using Bruker Topsin. Deuterated solvents used were obtained from Cambridge Isotope Laboratories. All NMR spectroscopy was performed at 300 K unless indicated otherwise. Coupling constants *J* are reported in Hz and rounded to the nearest 0.1 Hz. All NMR peaks are reported in parts per million relative to either TMS or residual solvent. Low-resolution mass spectrometry (LRMS) was performed on a Bruker amazon SL quadrupole ion trap mass spectrometer using electrospray ionisation (ESI). High-resolution mass spectrometry (HRMS)was performed on a Thermo Velos Pro Orbitrap mass spectrometer via syringe infusion on the ionMax Electrospray Ionisation source. Fourier transformed infrared spectroscopy (FTIR) was performed on a Bruker Platinum Alpha-E FTIR. Samples were analysed neat and signals reported with frequency of maximum absorbance (vmax/cm-1). Melting points were recorded on a Standford Research Systems OptiMelt at 3 °C per min.

#### Synthesis and Characterisation Data for Lead Actives C16–C18

Lead active compounds **C16**–**C18** were prepared according to the route shown in Scheme 1.

*1,4,8,11-tetraazatricyclo[9.3.1.14,8]hexadecane*

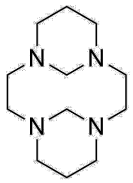


To cyclam (0.954 g, 4.76 mmol) in water (60 mL) was added formaldehyde (37% in water, 0.90 mL, 11 mmol) at 0°C. The mixture was stirred for 3 h, resulting in formation of a white precipitate. The mixture was filtered under reduced pressure and the residue was washed with water (50 mL) and dried in vacuo to give a white solid 0.893 g, 84%). **M.P.** 106–108 ºC (lit = 106–108 ºC). **$^{1}$H NMR** (300 MHz, $CDCl_3$): 1.18 (d, 2H, *J* 13.3), 2.19–2.46 (m, 6H), 2.53–2.71 (m, 4H), 2.76–2.94 (m, 6), 3.15 (d, 4H, 9.66), 5.43 (d, 2H, *J* 10.8). **LRMS** (ESI+): *m/z* 224.91 $[M+H]^+$. Data in agreement with literature. [1]

*Di-tert-butyl 4,11-di(prop-2-yn-1-yl)-1,4,8,11-tetraazacyclotetradecane-1,8-dicarboxylate* ***2***

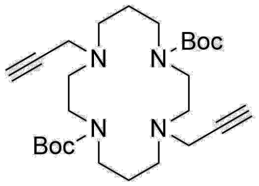


To a solution of 1,4,8,11-tetraazatricyclo[9.3.1.14,8]hexadecane (0.983 g, 4.39 mmol) in acetonitrile (60 mL) was added propargyl bromide (80% in toluene, 1.6 mL, 10.8 mmol) and the solution stirred at rt overnight. The mixture was concentrated under reduced pressure to dryness before being suspended in MeOH (40 mL). Aqueous NaOH solution (2.5 M, 60 mL) was added and the mixture was stirred at rt for 2 hours. The pH was adjusted to 11 before the addition of Boc anhydride (1.5 mL, 6.88 mmol) and the mixture stirred at rt overnight. The product was isolated via vacuum filtration and washed with water (60 mL) and dried in vacuo to leave the product as an off white solid (1.632 g, 81%). **MP**: 149–152 °C (lit. 153–155 °C). **$^{1}$H**

**NMR** (300 MHz, $CDCl_3$): 1.46 (s, 18H), 1.65–1.78 (m, 4H), 2.46–2.58 (m, 4H), 2.59–2.72 (m, 4H), 3.21–3.48 (m, 12H). **LRMS** (ESI+): *m/z* 477.36 $[M+H]^+$. Data in agreement with literature. [1]

*10-(3-(4-(2-Azidoethyl)piperazin-1-yl)propyl)-2-chloro-10H-phenothiazine* ***4***

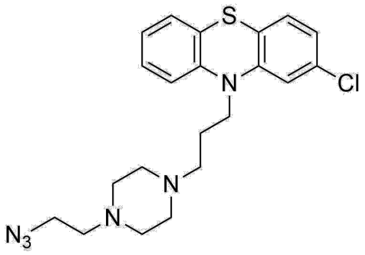


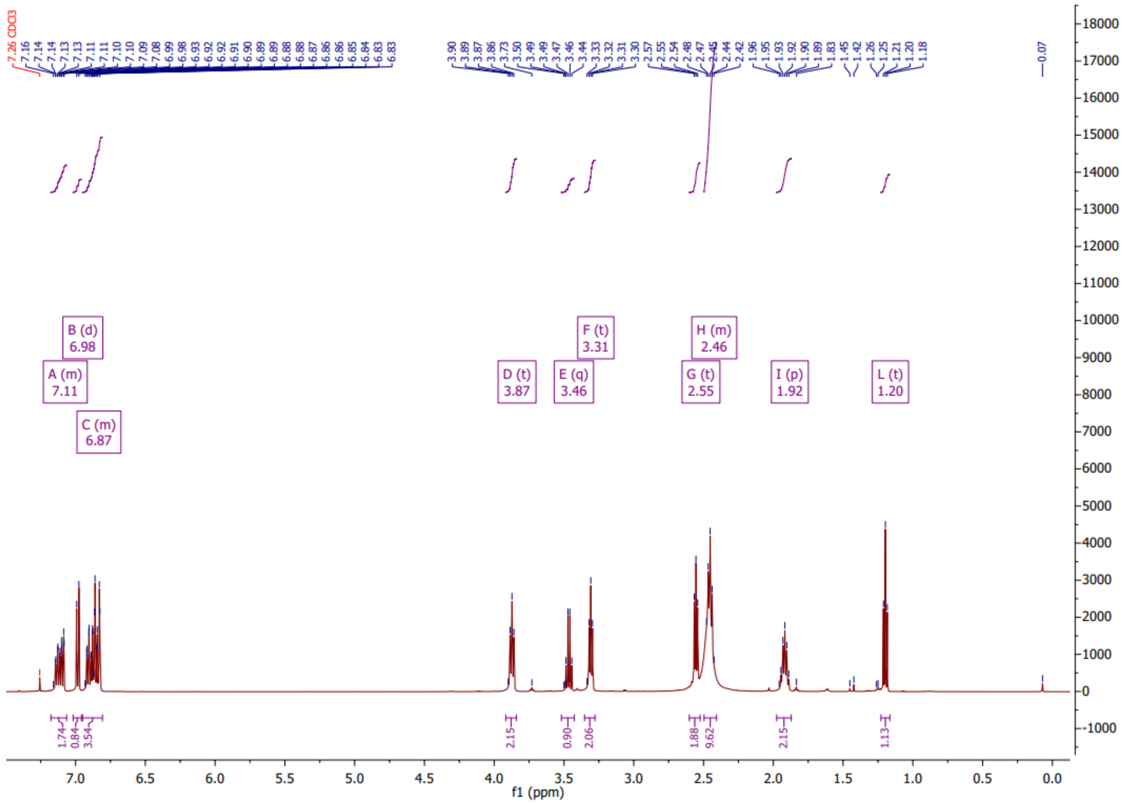

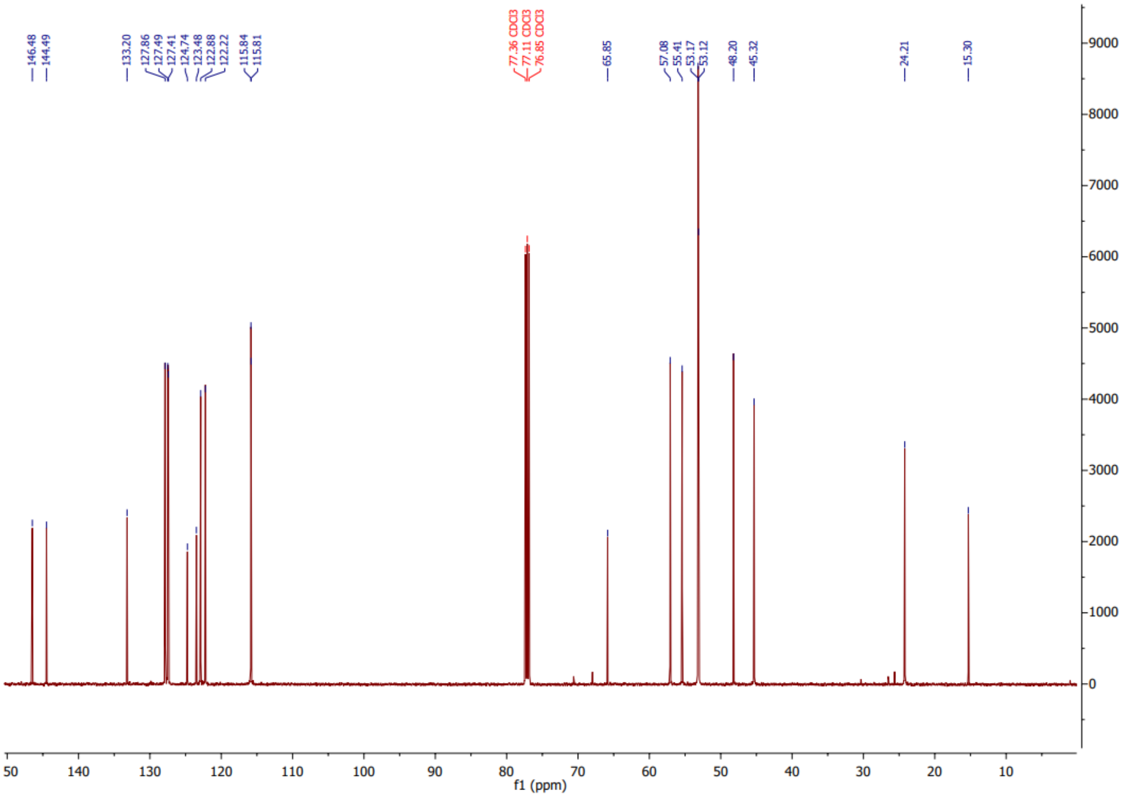


To a solution of perphenazine **3** (0.993 g, 2.46 mmol) in anhydrous DMF (10 mL) was added sodium azide (0.581 g, 8.93 mmol), DPPA (1.1 mL, 5.1 mmol) and DBU (0.75 mL, 5.02 mmol) and the vessel backfilled with $N_2$. The mixture was heated to 70 ºC and stirred for 24 h. Water (25 mL) was added dropwise and the product extracted with DCM (8 x 50 mL). The combined extracts were washed with water (4 x 200 mL), brine (2 x 200 mL) and dried over magnesium sulfate before being concentrated under reduced pressure. The crude product was purified via reversed phase chromatography (gradient of 5% MeOH in water to 100% MeOH) to leave the product as a red oil (0.832 g, 79%). **$^{1}$H NMR** (500 MHz, $CDCl_3$): 1.19 (qn, 2H, *J* 6.9), 2.39–2.53 (m, 8H), 2.55 (t, 2H, *J* 6.3), 3.31 (t, 2H, *J* 6.8), 3.87 (t, 2H, *J* 6.7), 6.84–6.93 (m, 4H), 6.98 (d, 1H, *J* 8.1), 7.06–7.16 (m, 2H). **$^{13}$C NMR** (125 MHz, $CDCl_3$): 24.21, 45.32, 48.20, 53.17, 55.40, 57.08, 115.81, 122.22, 122.87, 123.48, 124.74, 127.41, 127.49, 127.86, 133.20, 144.49, 146.47. **LRMS** (ESI+): *m/z* 429.19 $[M+H]^+$. **HRMS** (ESI+): *m/z* calc. for $C_{21}H_{26}Cl_1N_6S_1$ 429.16227 $[M+H]^+$, found 429.16195. **FTIR** (ATR) $\nu_{max}$/cm$^{-1}$: 2940, 2809, 2097, 1566, 1455, 1407, 1279, 1242, 1158, 1140, 1033.

*Di-tert-butyl 4,11-bis((1-(2-(4-(3-(2-chloro-10H-phenothiazin-10-yl)propyl)piperazin-1-yl)ethyl)-1H-1,2,3-triazol-4-yl)methyl)-1,4,8,11-tetraazacyclotetradecane-1,8-dicarboxylate* ***5***

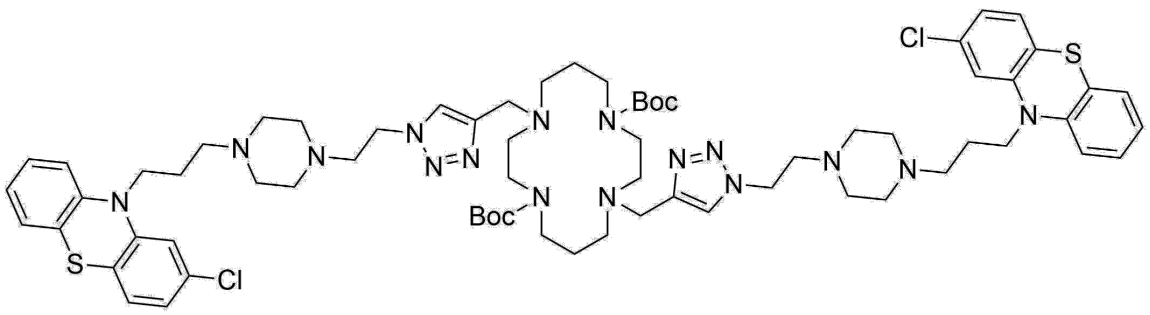


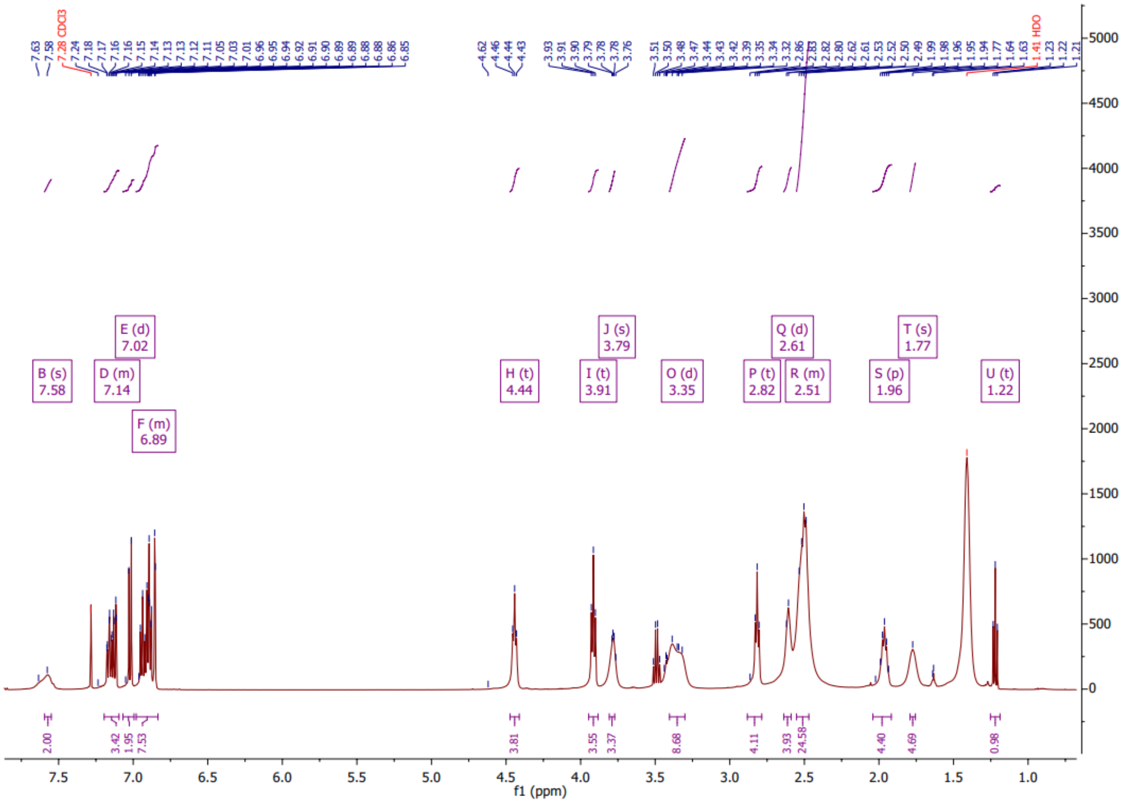

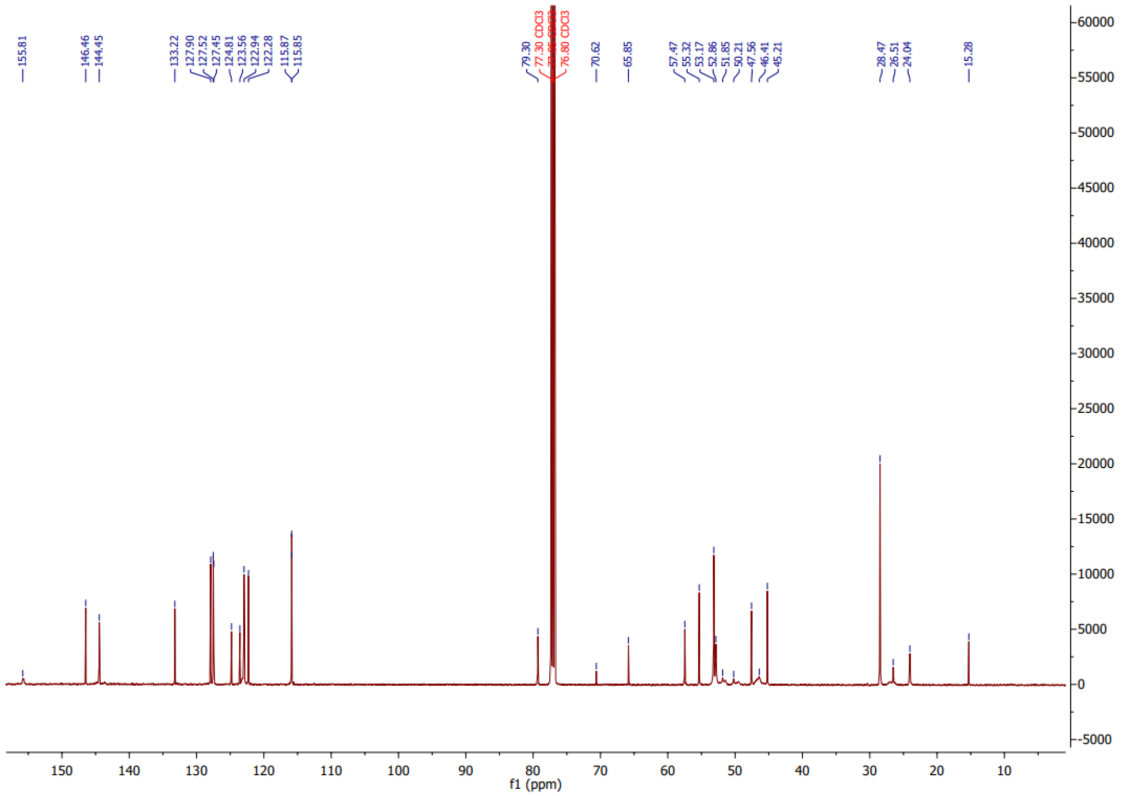

A solution of **2** (0.146 g, 0.31 mmol), copper sulfate pentahydrate (15.6 mg, 0.06 mmol), and sodium ascorbate (32.7 mg, 0.16 mmol) in THF/water (6:9, 15 mL) was flushed with argon and stirred at rt for 5 min. A solution of **4** (0.255 g, 0.59 mmol) in THF (1 mL) was added and the mixture stirred at rt overnight. Ammonium chloride (sat. aqueous, 15 mL) was added dropwise and the product extracted with DCM (2 x 50 mL) to leave the crude product as an orange oil. The mixture was purified via reversed phase chromatography (5% MeOH/water ramping to 100% MeOH) to leave the product as a white powder (0.188 g, 46%). **MP**: 84–88 ºC. **$^1$H NMR** (500 MHz, $CDCl_3$): 1.41 (s, 18H), 1.77 (bs, 4H), 1.90–2.03 (m, 4H), 2.36–2.64 (m, 28H), 2.77–2.89 (m, 4H), 3.25–3.45 (m, 8H), 3.77 (bs, 4H), 3.86–3.96 (m, 4H), 4.40–4.52 (m, 4H), 6.84–7.22 (m, 14H), 7.56 (bs, 2H). **$^{13}$C NMR** (125 MHz, $CDCl_3$): 146.46, 144.45, 133.22, 127.90, 127.52, 127.45, 124.81, 123.56, 122.94, 122.28, 115.87, 79.29, 70.62, 65.84, 57.47, 55.32, 53.16, 52.85, 47.56, 45.21, 28.47, 26.51, 24.04, 15.27. **LRMS** (ESI+): *m/z* 1335.76 $[M+H]^+$. **HRMS** (ESI+): *m/z* Calc. for $C_{68}H_{96}Cl_2N_{16}O_4S_2$ 667.33040 $[M+2H]^{2+}$, found 667.32932. **FTIR** (ATR) $\nu_{max}$/cm$^{-1}$: 2936, 2808 1682, 1566, 1456, 1408, 1364, 1245, 1127, 1110.

*1,8-Bis((1-(2-(4-(3-(2-chloro-10H-phenothiazin-10-yl)propyl)piperazin-1-yl)ethyl)-1H-1,2,3-triazol-4-yl)methyl)-1,4,8,11-tetraazacyclotetradecane **C16***

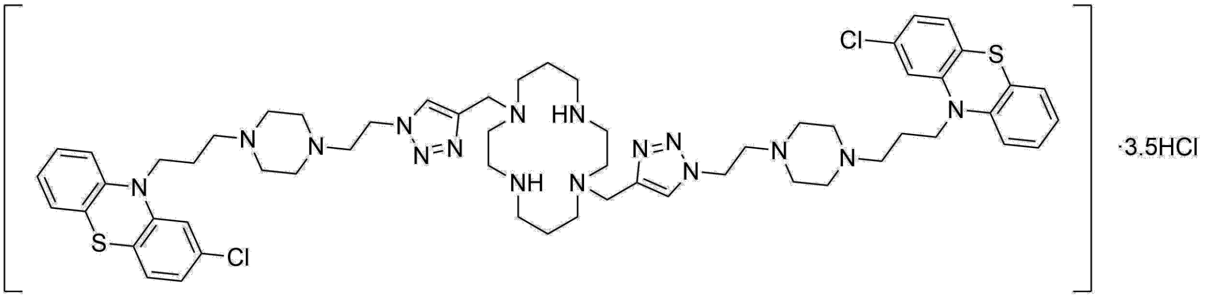

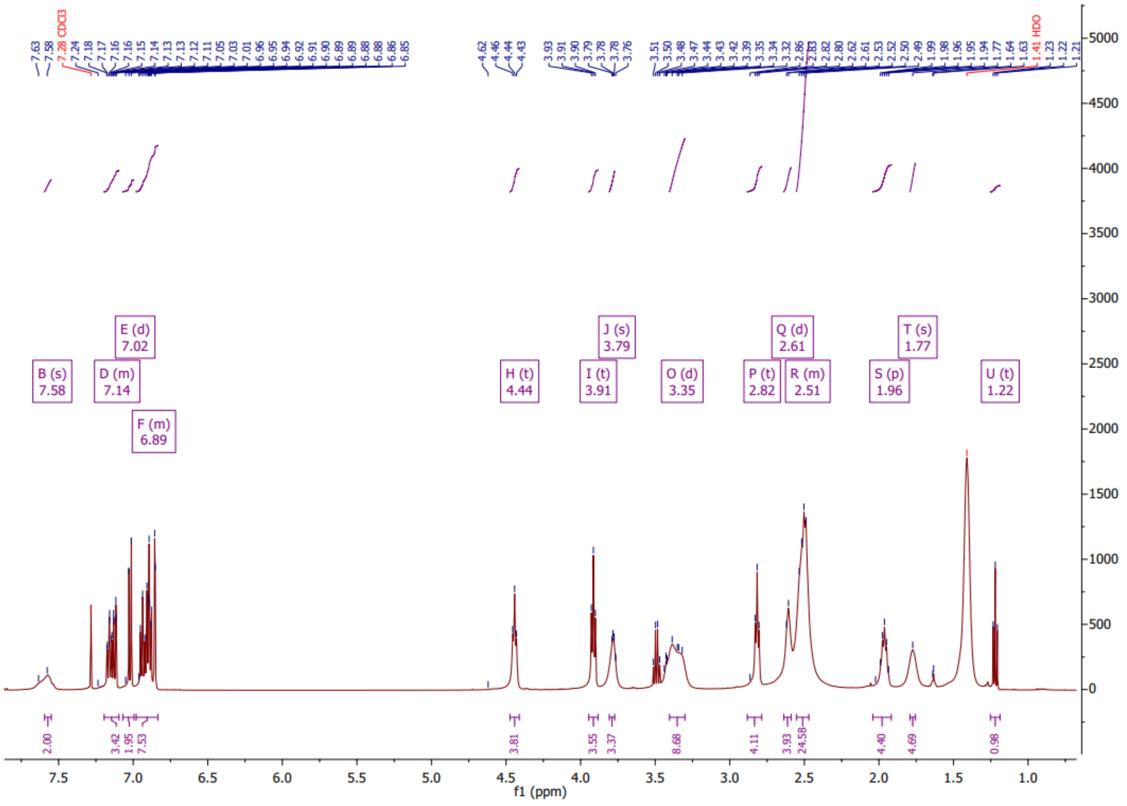

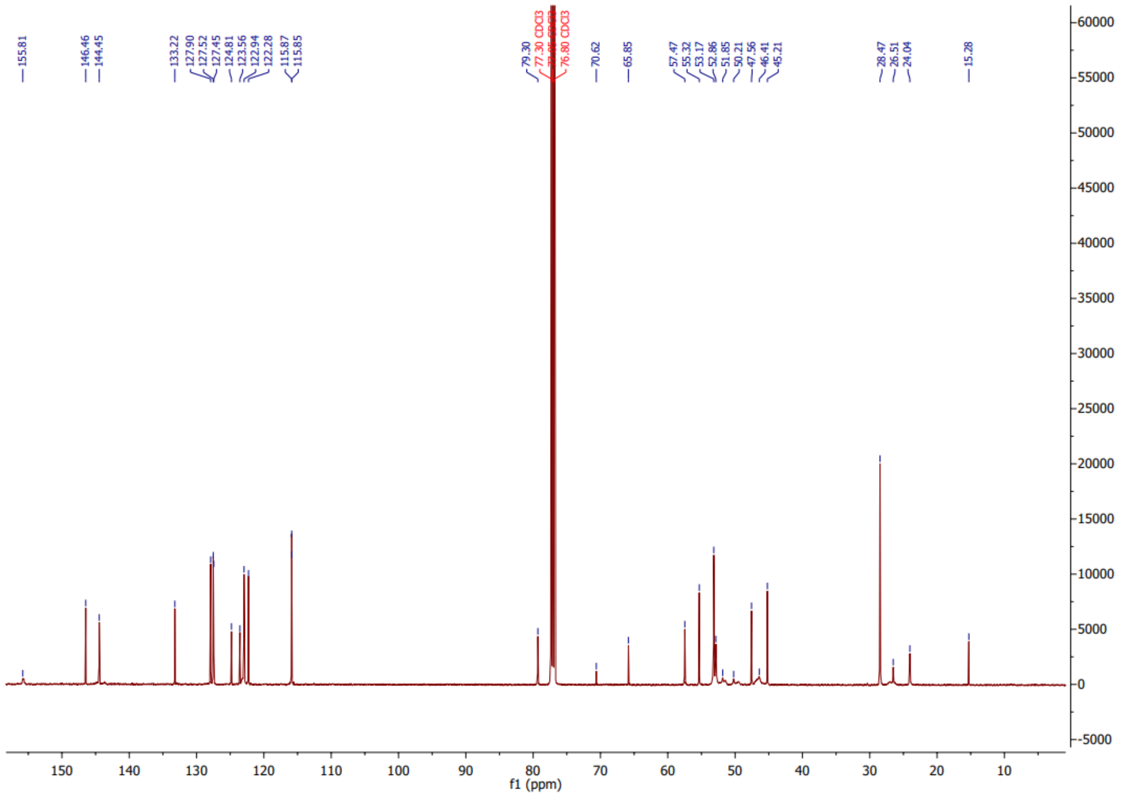

To a solution of Boc-protected bis-perphenazine cyclam **4** (196 mg, 0.173 mmol) in dioxane (5 mL) and water (0.2 mL) was added HCl (4 M in dioxane, 1.8 mL, 7.8 mmol) at 0 ºC under nitrogen. The mixture was stirred at rt for 1 h before being concentrated under reduced pressure. The product was purified via flash column chromatography (reversed phase, 5% MeOH in water to 100% MeOH) to yield the product as an off-white powder (160 mg, 74%). **$^{1}$H-NMR** (200 MHz, DMSO, 350K): 1.64–1.89 (m, 10H), 2.26–2.45 (m, 24H), 2.56–2.65 (m, 4H) 2.67–2.85 (m, 12H,), 3.74 (s, 4H), 3.88–4.01 (m, 4H), 4.36–4.51 (m, 4H), 6.94–7.29 (m, 14H), 7.95 (s, 2H). **$^{13}$C NMR**: 145.90, 143.17, 142.25, 133.0, 128.20, 127.92, 127.65, 125.74, 124.23, 123.59, 123.05, 122.50, 116.58, 115.67, 60.31 55.29, 54.05, 52.99, 50.17, 48.52, 44.83, 43.26, 21.80, 21.00. **LRMS** (ESI+): *m/z* 113.62. **HRMS** (ESI+): *m/z* calc. for $C_{58}H_{78}Cl_2N_{16}S_2$ 1133.54866 free base $[M+H]^+$, found 1133.54823. **FTIR** (ATR) $\nu_{max}/cm^{-1}$: 2968, 2801, 2186, 1682, 1566, 1455, 1408, 1244, 1151, 1050.

*1,8-Bis((1-(2-(4-(3-(2-chloro-10H-phenothiazin-10-yl)propyl)piperazin-1-yl)ethyl)-1H-1,2,3-triazol-4-yl)methyl)-1,4,8,11-tetraazacyclotetradecane, $Zn(ClO_4)_2$ complex **C17**.*

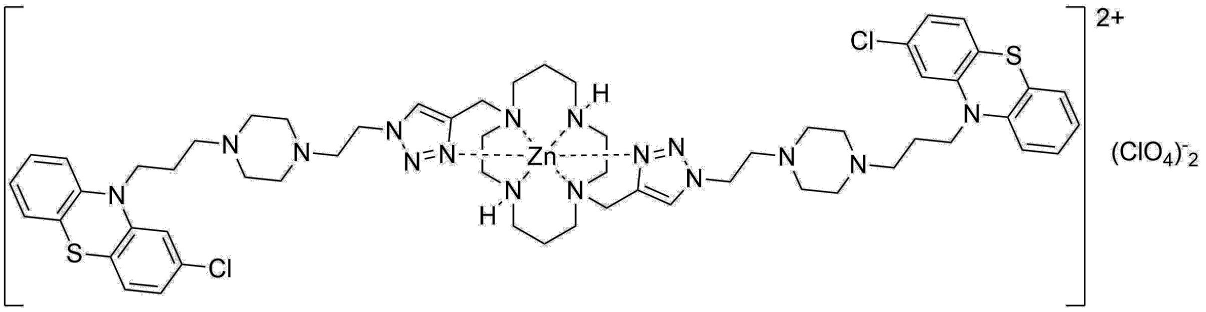

A solution of **C16** (44.8 mg, 0.04 mmol) in methanol (2 mL) was added to a suspension of Ambersep 900 hydroxide form resin (~150 mg) in methanol (15 mL) that had been pre-swollen for 30 min. The mixture was stirred at rt for 15 min and filtered. Zinc perchlorate hexahydrate (16.5 mg, 0.04 mmol) was added to the filtrate and the mixture stirred at reflux for 16 h. The solution was concentrated under reduced pressure and washed with diethyl ether (5 mL) to yield the product as a white solid (31.8 mg, 63%). **LRMS** (ESI+): m/z 1299.49 $[M–ClO_4]^+$. **HRMS** (ESI+): m/z Calc. for C58H78Cl3N16S2O4Zn 1297.41660 $[M–ClO_4]^+$, found 1297.41746. **FTIR** (ATR) $\nu_{max}/cm^{-1}$: 2927, 2870, 2815, 1564, 1455, 1072, 789, 750, 622. 89, 750, 622.

*1,8-Bis((1-(2-(4-(3-(2-chloro-10H-phenothiazin-10-yl)propyl)piperazin-1-yl)ethyl)-1H-1,2,3-triazol-4-yl)methyl)-1,4,8,11-tetraazacyclotetradecane, $Cu(ClO_4)_2$ complex* ***C18***.

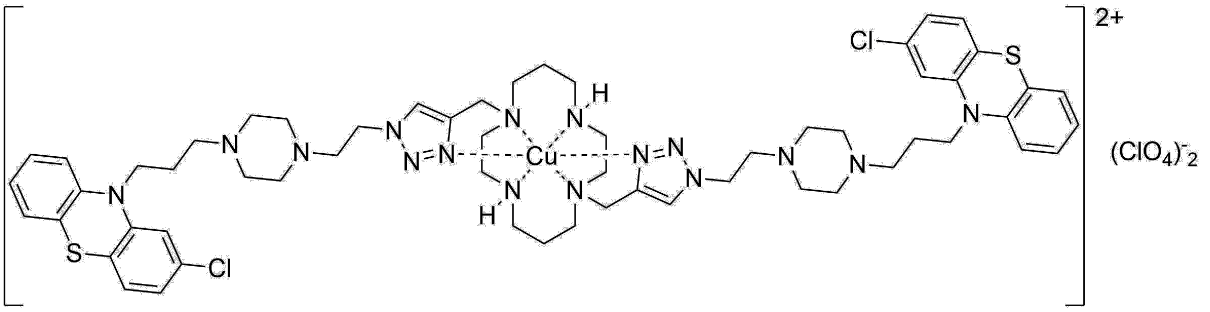

A solution of **C16** (39.6 mg, 0.035 mmol) in methanol (2 mL) was added to a suspension of Ambersep 900 hydroxide form resin (~150 mg) in methanol (15 mL) that had been pre-swollen for 30 min. The mixture was stirred at rt for 15 min and filtered. Copper perchlorate hexahydrate (13.2 mg, 0.035 mmol) was added to the filtrate and the mixture stirred at rt for 16 h. The solution was concentrated under reduced pressure and washed with diethyl ether (5 mL) to yield the product as a blue solid (28.3 mg, 64%). **LRMS** (ESI+): m/z 1296.51$[M–ClO_4]^+$. **HRMS** (ESI+): m/z Calc. for C58H78Cl3N16O4Cu 1296.41747 $[M–ClO_4]^+$, found 1296.41740. **FTIR** (ATR) $\nu_{max}/cm^{-1}$: 2937, 2814, 1565, 1455, 1082, 921, 747, 621.

**Method for screening effects of compounds against synthetic Aβ42**

Aβ42 (0.25 mg, Genscript) was solubilised in aqueous NaOH solution (60 mM, 62.5 µL) and incubated for 5 min at room temperature. The solution was diluted into deionised water (175 µL) and bath sonicated for 5 min. 10X PBS (25 µL, pH 7.0) was added and the solution centrifuged at 14000 *g* for 10 min. The supernatant was extracted and concentration measured via spectrophotometer ($\varepsilon_{214}$ = 95452). Peptide samples were prepared in triplicate in 96-well plates in aqueous $NaH_2PO_4$ buffer (25 mM) with NaCl (150 mM) at pH 7.4, and ThT (40 µM) to a final volume of 100 µL. Aβ42 (10 µM) was incubated alone or in the presence of 2 µM compound (Fig. S1B). Samples were incubated at 37 °C in a POLARstar Omega microplate reader (BMG Labtech), with excitation at 440 nm and the fluorescence emission recorded at 480 nm with double orbital shaking at 700 rpm for ~15 h.

**Method for screening effects of compounds against recombinant Aβ40 and Aβ42**

Recombinant Aβ40 and Aβ42 was expressed and purified as outlined in the main methods section, omitting the final SEC purification step. Concentrated lypophilised Aβ40 or Aβ42 were resuspended in aqueous $NaH_2PO_4$ buffer (25 mM) with NaCl (150 mM) pH 7.4 and concentration was measured via spectrophotometer ($\varepsilon_{275}$ = 1450). Peptide samples were prepared in triplicate in 96-well plates in $NaH_2PO_4$ (25 mM) with NaCl (150 mM) at pH 7.4, and ThT (40 μM) to a final volume of 100 μL. Aβ40 or Aβ42 (10 μM) was incubated alone or in the presence of compound (25 μM, Fig. S1C). Samples were incubated at 37 °C in a POLARstar Omega microplate reader (BMG Labtech), with excitation at 440 nm and the fluorescence emission recorded at 480 nm with double orbital shaking at 700 rpm for >15 h.

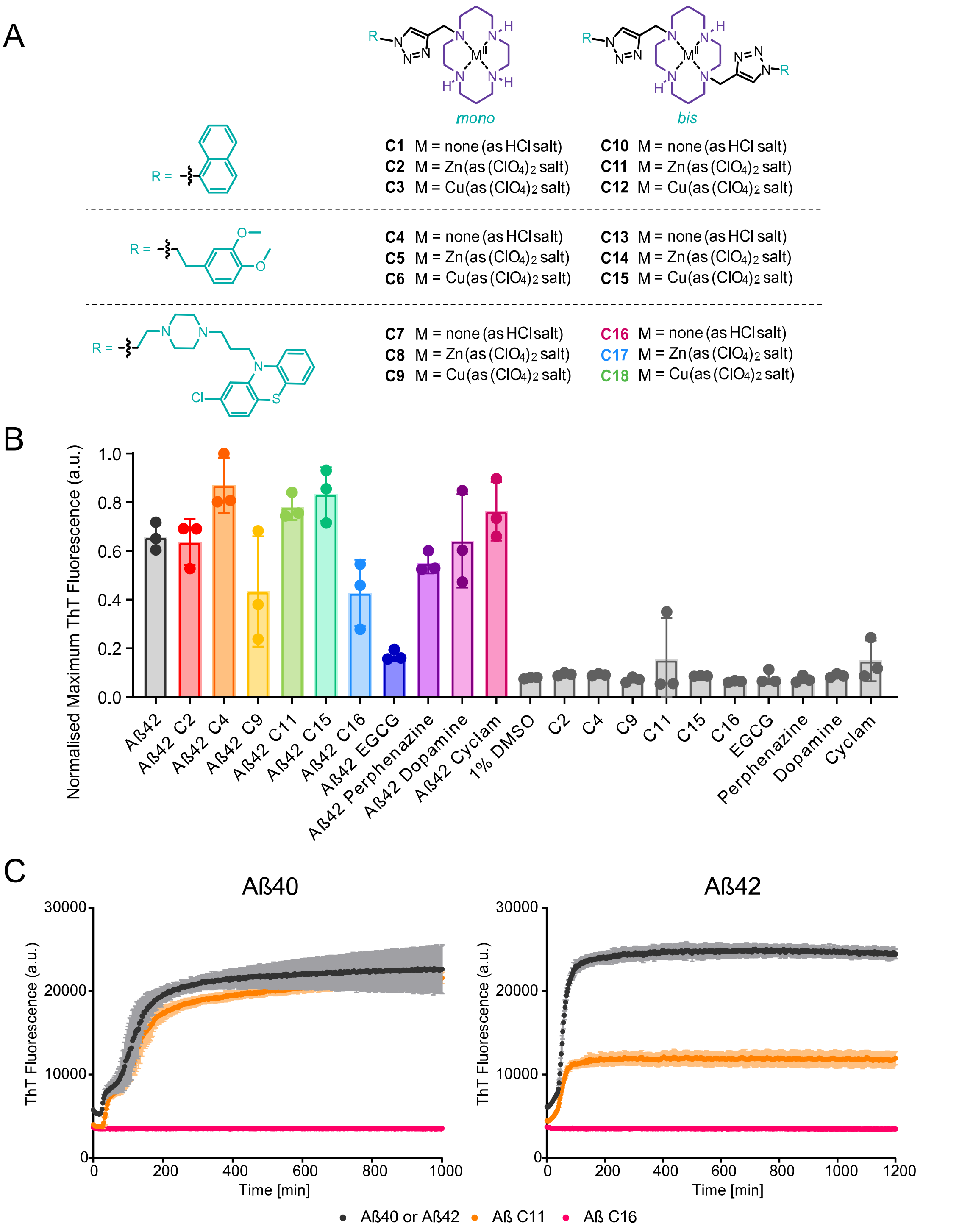


**Fig. S1.** *(A)* The compounds used in this study combine a cyclam macrocycle (purple) with pendant moieties (blue), conjugated via a click-derived triazole, and deployed either as the unmetallated HCl salt, the zinc perchlorate complex, or the copper(**II**) perchlorate complex. The pendant groups are derived from either a naphthyl (**C1**–**C3**, **C10**–**C12**), dopamine-derived (**C4**–**C6**, **C13**–**C15**), or pherphenazine-derived (**C7**–**C9**, **C16**–**C18**) moiety, and conjugates are either 'single-armed' compounds that contain only a single triazolyl-pendant group (**C1**–**C9**) or 'double-armed' structures (**C10**–**C18**). *(B)* Preliminary compound screening against 10 μM synthetic monomeric Aβ42 using 2 μM compound. Samples were measured in triplicate under shaking conditions at 37 °C. Data were corrected for time zero and normalised as a function of the maximum fluorescence intensity using GraphPad Prism. N = 4. *(C)* ThT assays using recombinant 10 μM Aβ40 or Aβ42 (black) and 25 μM of **C11** (orange) or **C16** (pink). Samples were measured in triplicate under shaking conditions at 37 °C. Error bars represent SD.

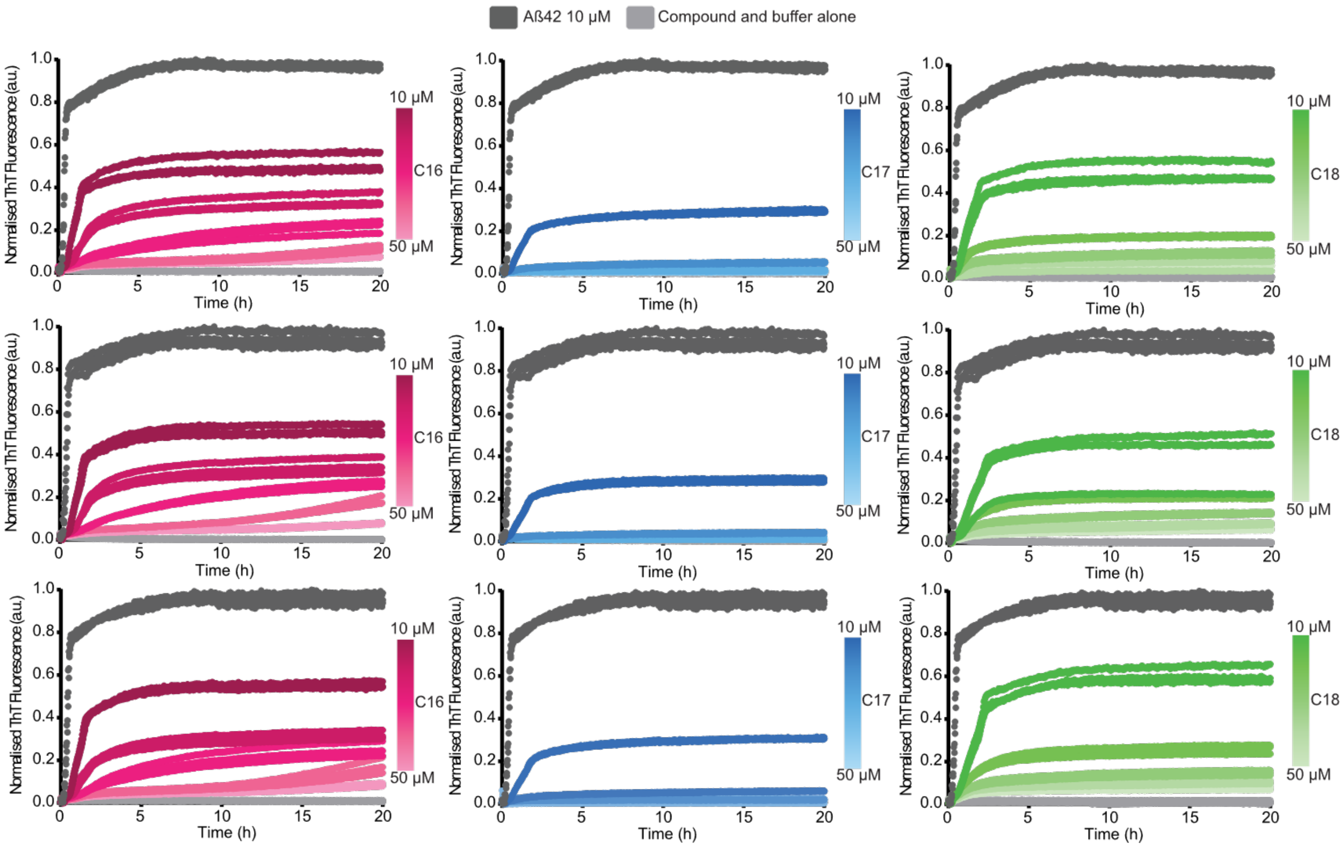


**Fig. S2.** Concentration dependence of the inhibition of Aβ42 aggregation by C16 (pink), C17 (blue) and C18 (green). 10 µM monomeric Aβ42 in the absence or presence of either 1% DMSO (dark grey) or 10–50 µM of compound. Samples were measured in triplicate under quiescent conditions at 37 °C. Data from three repeat experiments are shown, each using protein from separate purifications. Data were corrected for time zero and normalised as a function of the maximum fluorescence intensity using GraphPad Prism.

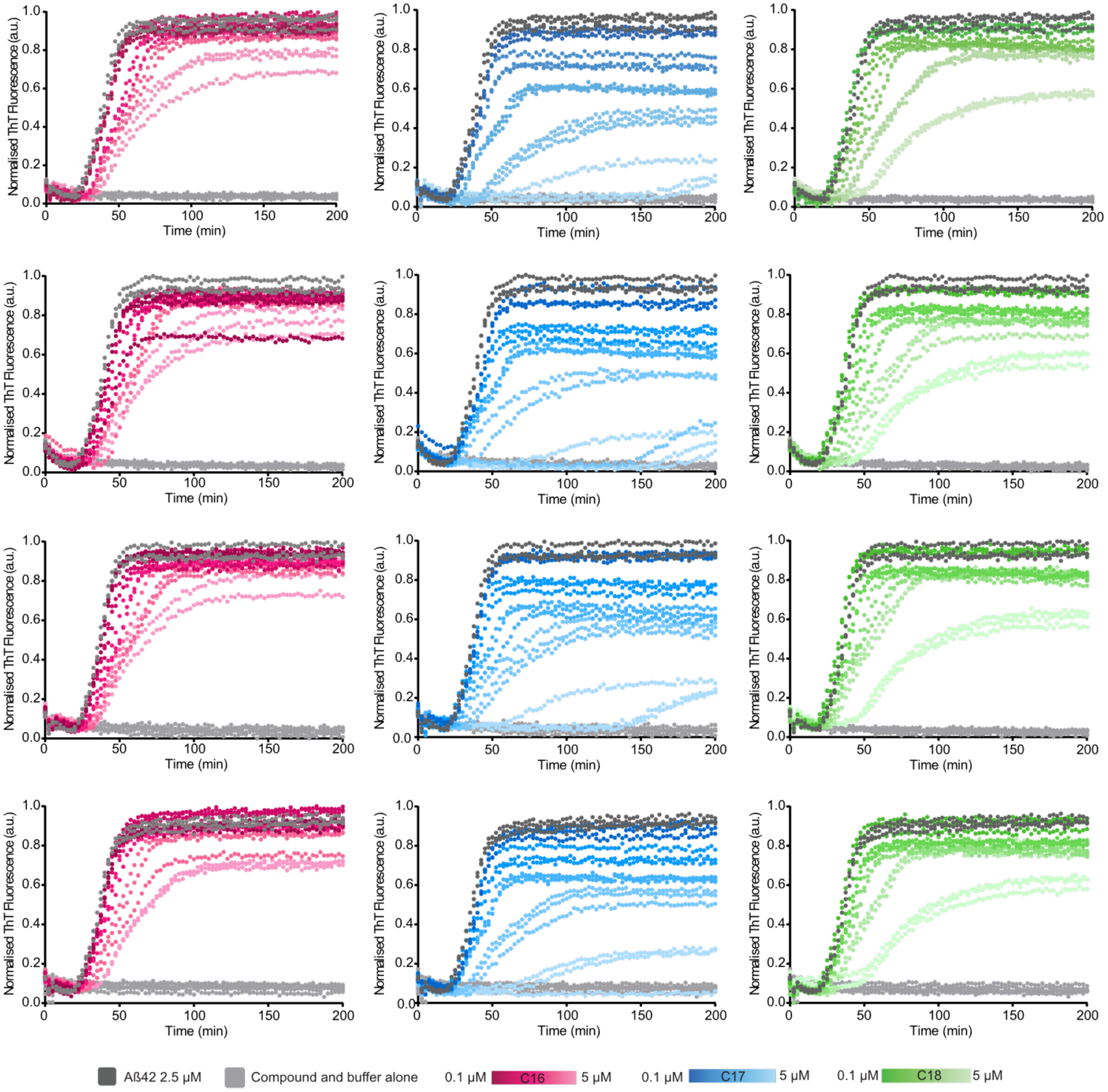


**Fig. S3.** Concentration dependence of the inhibition of Aβ42 aggregation by C16 (pink), C17 (blue) and C18 (green) over a lower range of inhibitor concentrations. 2.5 μM monomeric Aβ42 in the absence or presence of either 1% DMSO (dark grey) or 0.1, 1, 2, 3 or 5 μM of compound. Samples were measured in triplicate under quiescent conditions at 37 °C. Data from four repeat experiments are shown, each using protein from separate purifications. Data were corrected for time zero and normalised as a function of the maximum fluorescence intensity using GraphPad Prism.

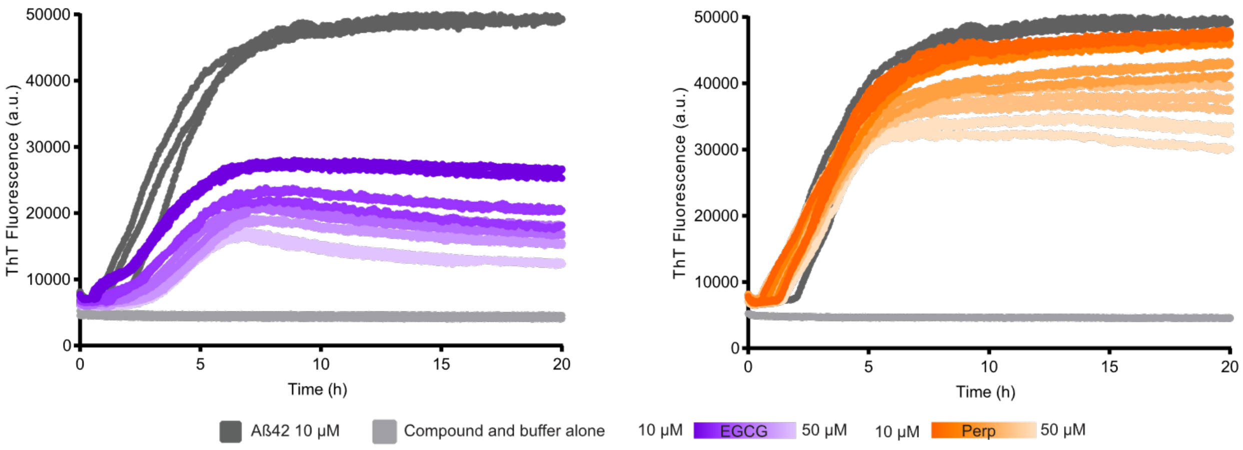


**Fig. S4.** Representative time-course showing concentration dependence of the inhibition of Aβ42 aggregation by EGCG (purple) and perphenazine (orange). 10 µM monomeric Aβ42 in the absence or presence of either 1% DMSO (dark grey) or 10–50 µM EGCG or perphenazine. Samples were measured in triplicate under quiescent conditions at 37 °C. Data were corrected for time zero and normalised as a function of the maximum fluorescence intensity using GraphPad Prism.

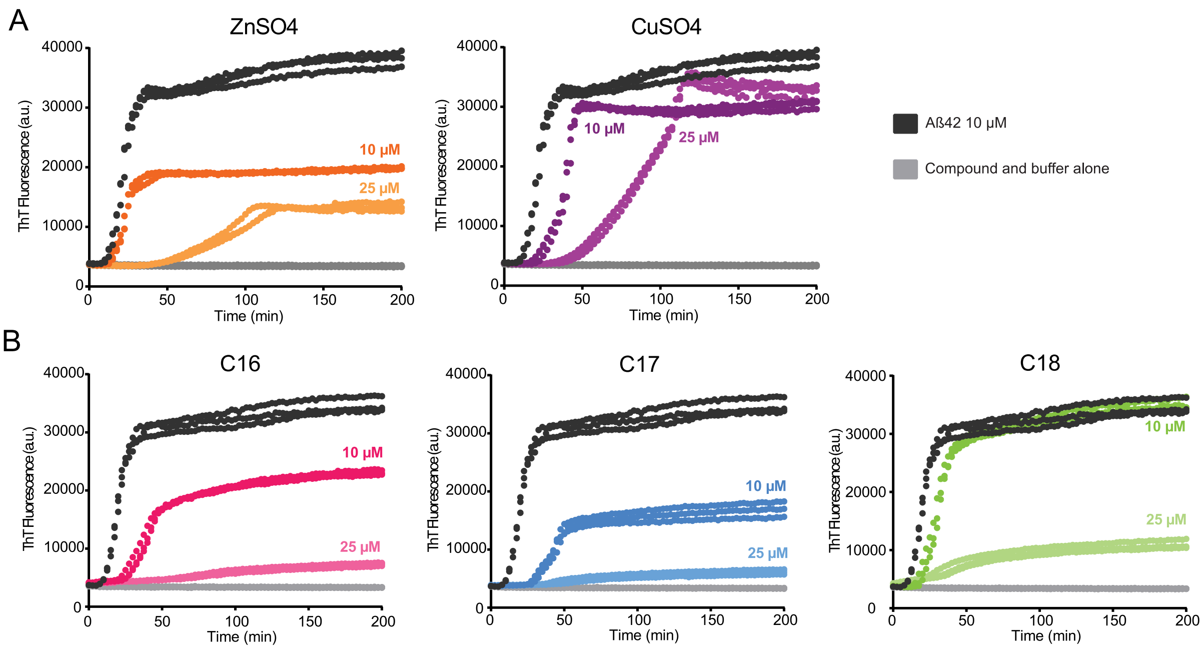


**Fig. S5.** *(A)* Representative time-courses showing concentration dependence of the inhibition of Aβ42 aggregation by Zn(**II**) (orange) and Cu(**II**) (purple). 10 µM monomeric Aβ42 in the absence or presence of either 1% DMSO (dark grey) or 10 µM or 25 µM of $ZnSO_4$ or $CuSO_4$. N = 2. *(B)* Representative time-courses showing inhibition of Aβ42 aggregation in the presence of EDTA. 10 µM monomeric Aβ42 in the absence or presence of either 1% DMSO (dark grey) or 10 µM or 25 µM of **C16** (pink), **C17** (blue) and **C18** (green). Buffer contained 200 µM EDTA. N =3. Samples were measured in triplicate under quiescent conditions at 37 °C. Data were corrected for time zero and normalised as a function of the maximum fluorescence intensity using GraphPad Prism.

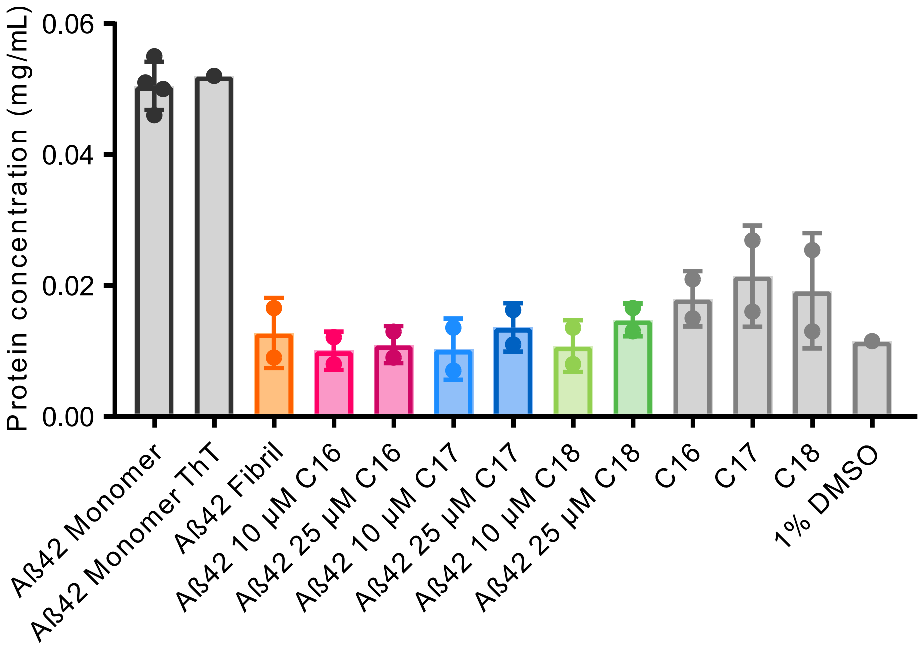


**Fig. S6.** Concentration of protein remaining in supernatant after Thioflavin T assay. 10 µM Aβ42 in the presence or absence of 1% DMSO (orange) or 10 µM or 25 µM of **C16** (pink), **C17** (blue) and **C18** (green). Monomeric 10 µM Aβ42, compounds alone and solvent used as controls. N = 2–4.

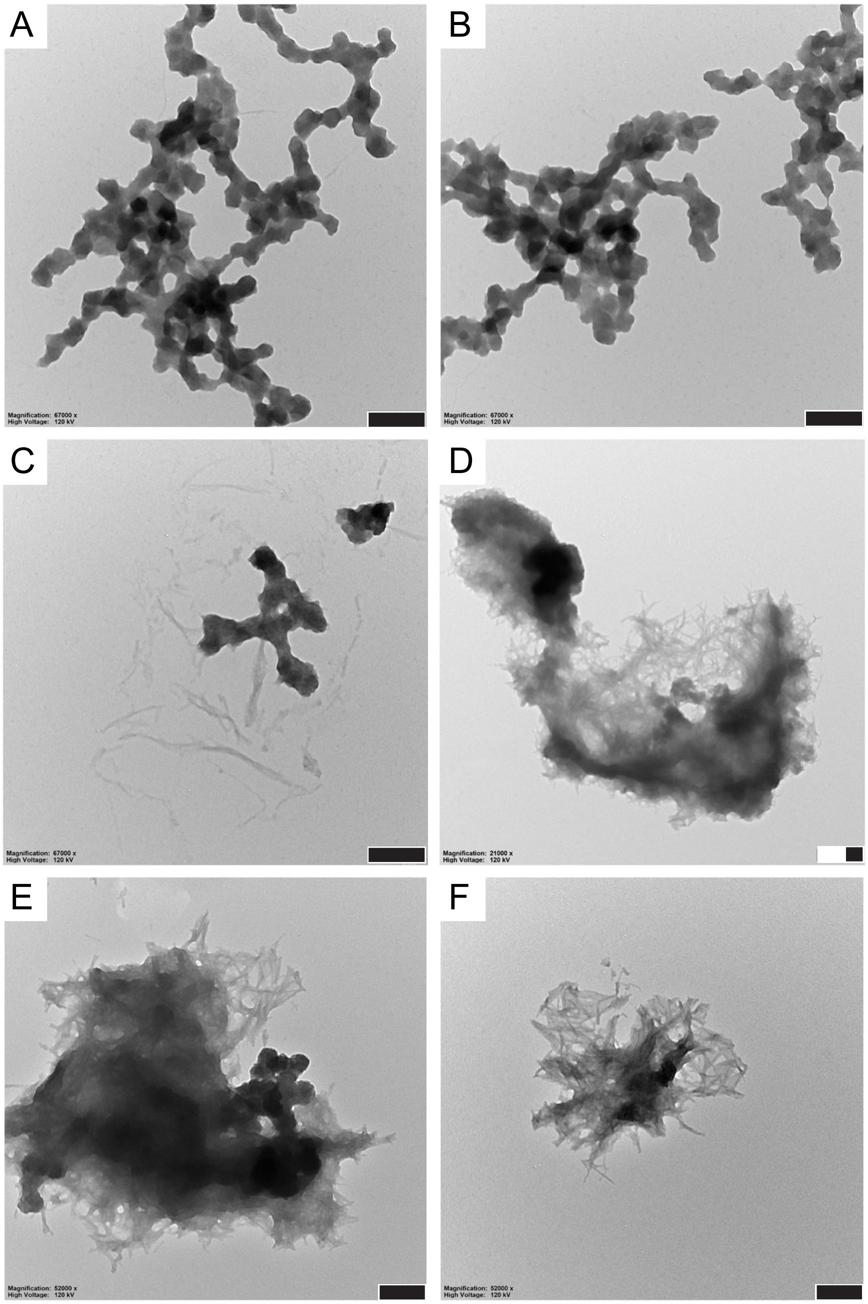


**Fig. S7.** Transmission electron microscopy images representative of the morphology of fibrils and aggregates formed by 10 µM Aβ42 when incubated in the presence of 10 µM **C16** (A,B), **C17** (C,D) and **C18** (E,F). Scale bar represents 200 nm.

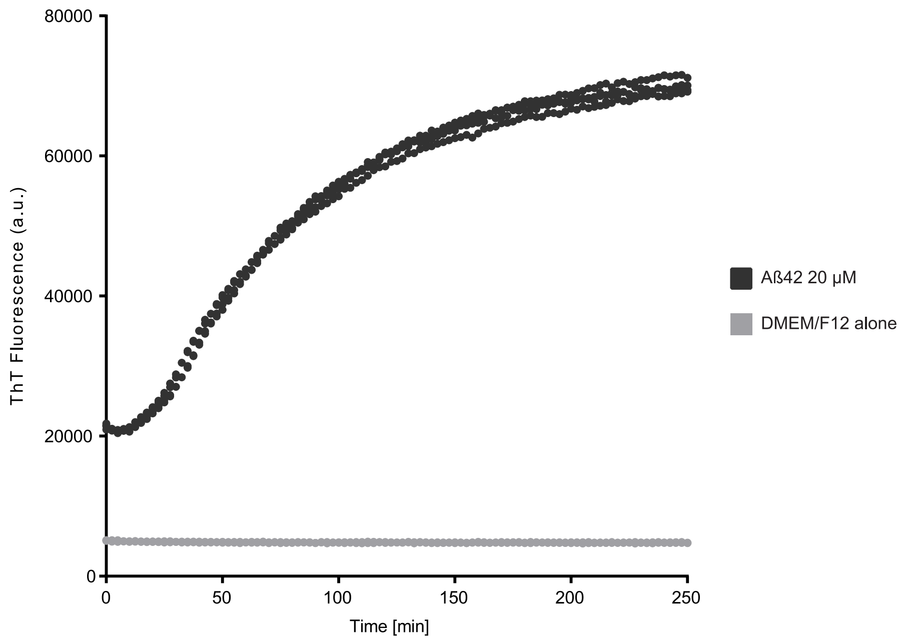


**Fig. S8.** Representative time-course of fibril formation by HFIP-treated 20 µM Aβ42 prepared for cell viability assay. DMEM/F12 cell media was used as buffer. Thioflavin T fluorescence reflects amyloid assembly. Samples were measured in triplicate under quiescent conditions at 37 °C.

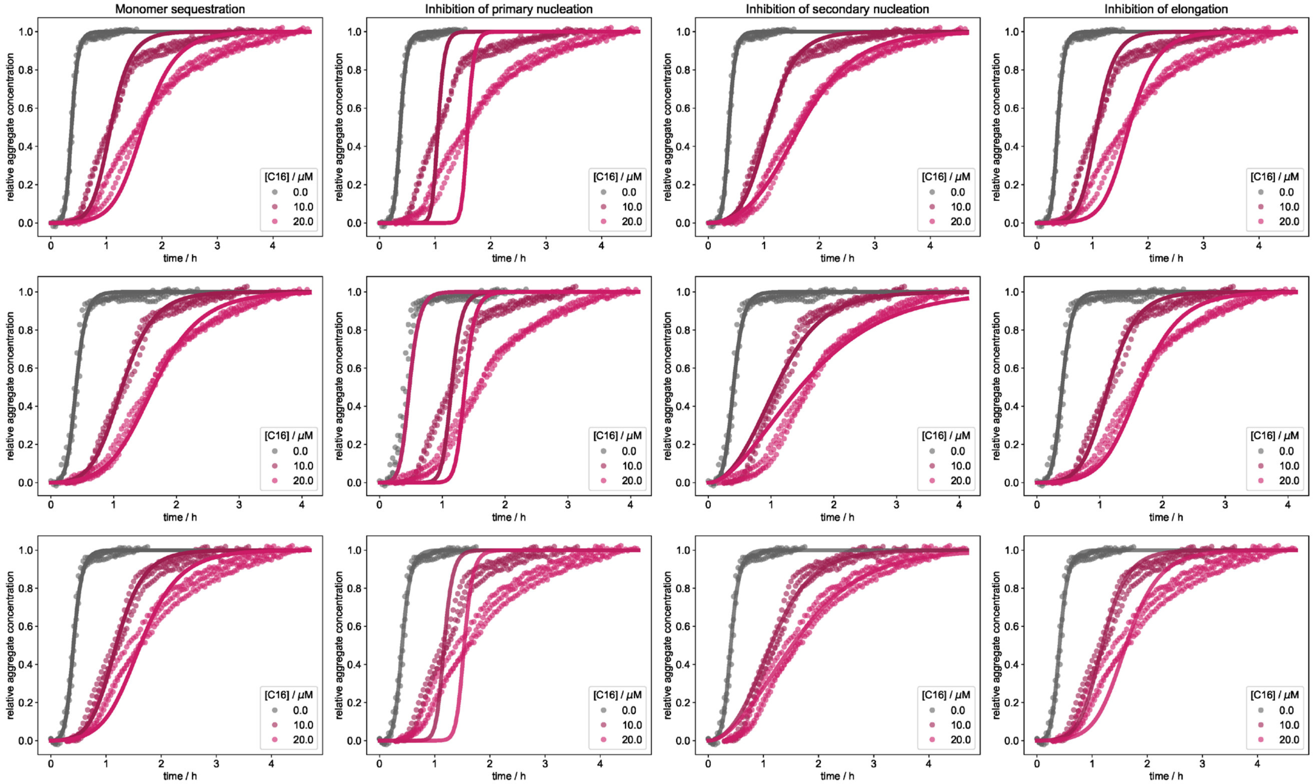


**Fig. S9.** Fitting results (solid lines) of the aggregation of 10 μM Aβ42 in the presence of **C16**, by means of the normalised kinetics measured in thioflavin T fluorescence assays. The fits are based on the kinetic parameters obtained from the unperturbed systems, allowing deviations in only the total monomer concentration, the rate constant of primary nucleation, secondary nucleation or elongation, respectively (columns left to right). Different rows show independent repeats of the experiment.

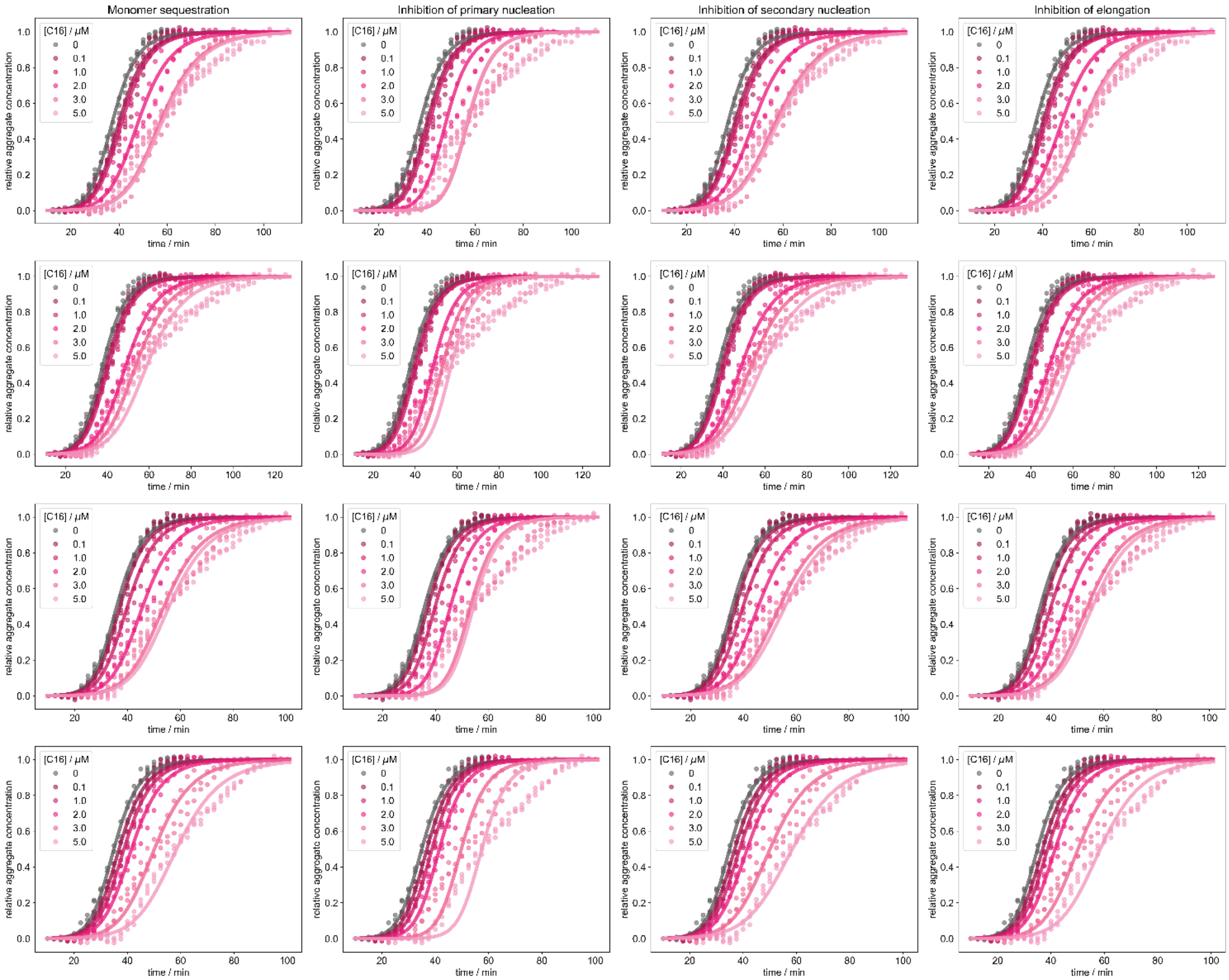


**Fig. S10.** Fitting results (solid lines) of the aggregation of 2.5 μM Aβ42 in the presence of **C16**, by means of the normalised kinetics measured in thioflavin T fluorescence assays. The fits are based on the kinetic parameters obtained from the unperturbed systems, allowing deviations in only the total monomer concentration, the rate constant of primary nucleation, secondary nucleation or elongation, respectively (columns left to right). Different rows show independent repeats of the experiment.

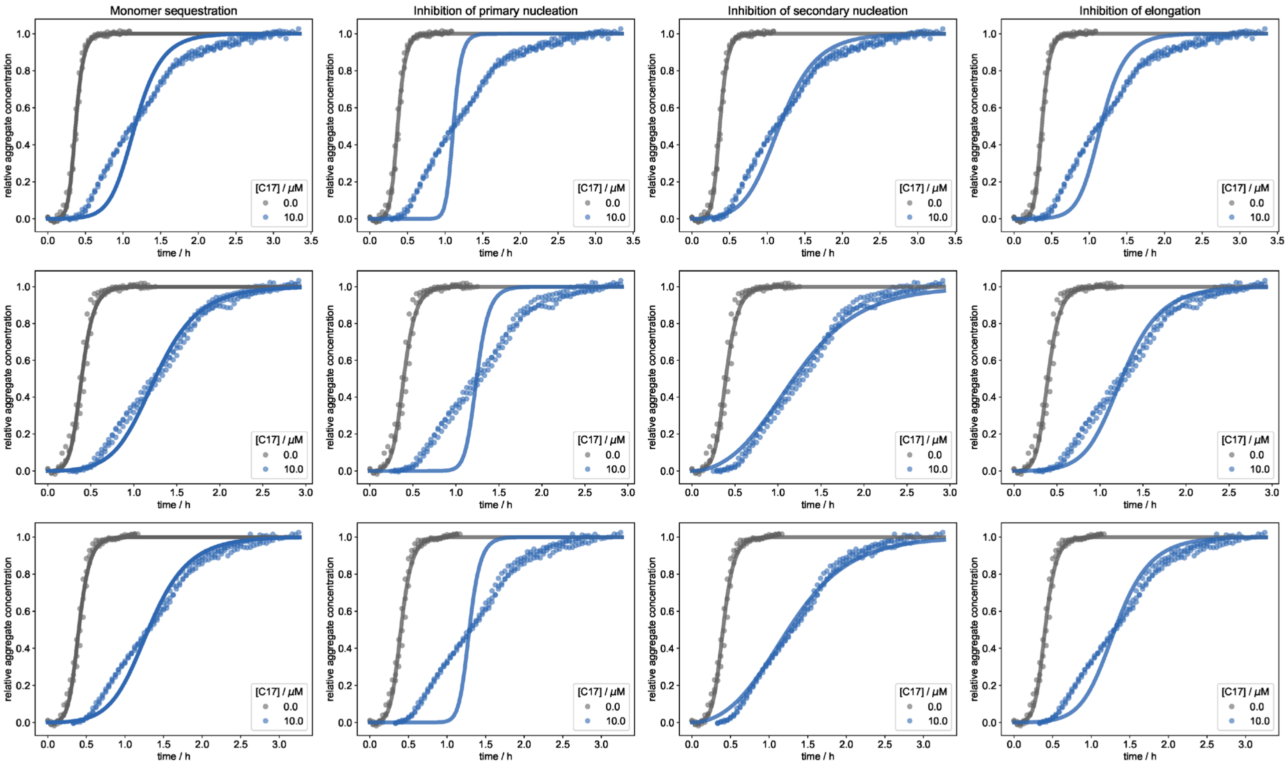


**Fig. S11.** Fitting results (solid lines) of the aggregation of 10 μM Aβ42 in the presence of **C17**, by means of the normalised kinetics measured in thioflavin T fluorescence assays. The fits are based on the kinetic parameters obtained from the unperturbed systems, allowing deviations in only the total monomer concentration, the rate constant of primary nucleation, secondary nucleation or elongation, respectively (columns left to right). Different rows show independent repeats of the experiment.

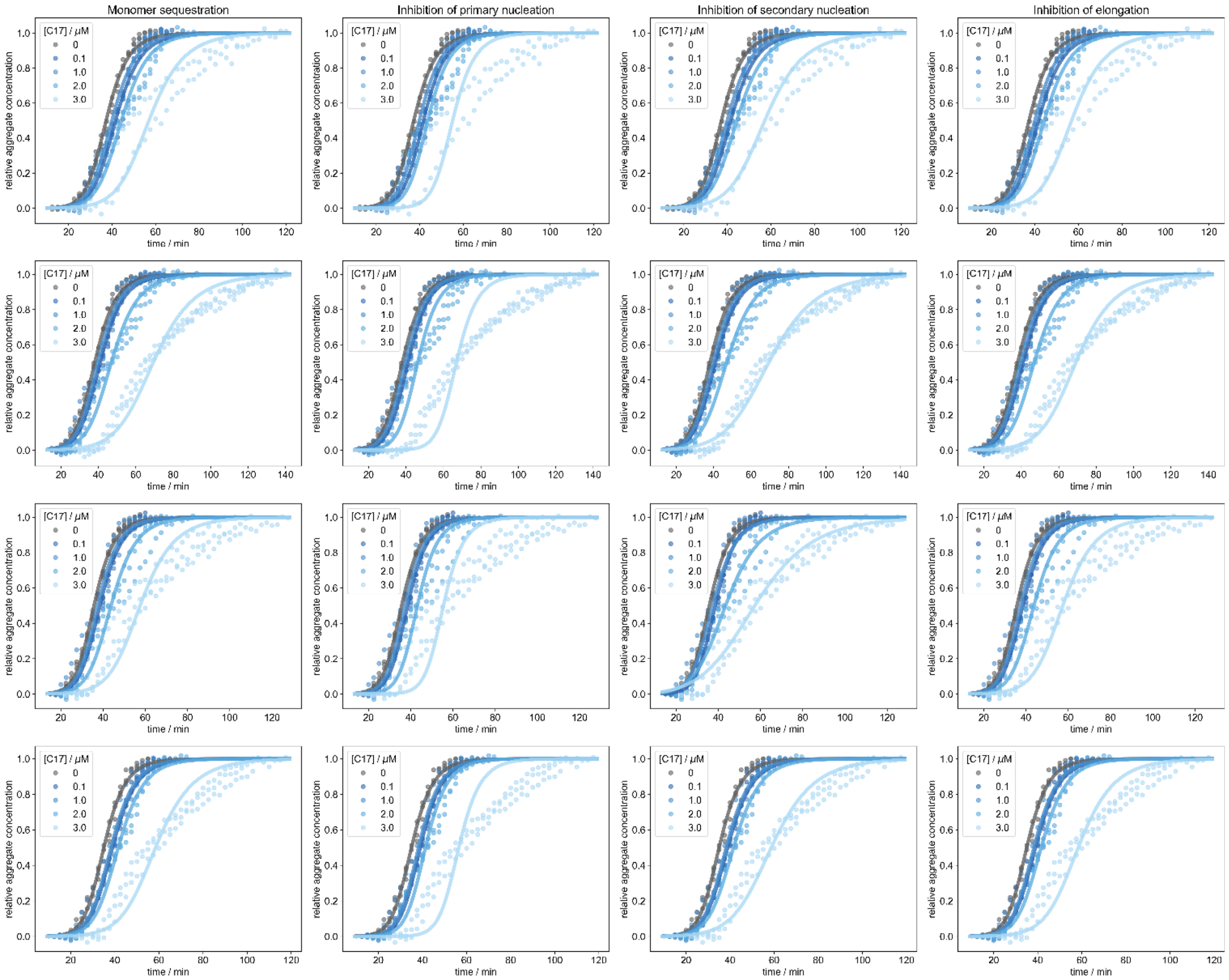


**Fig. S12.** Fitting results (solid lines) of the aggregation of 2.5 μM Aβ42 in the presence of **C17**, by means of the normalised kinetics measured in thioflavin T fluorescence assays. The fits are based on the kinetic parameters obtained from the unperturbed systems, allowing deviations in only the total monomer concentration, the rate constant of primary nucleation, secondary nucleation or elongation, respectively (columns left to right). Different rows show independent repeats of the experiment.

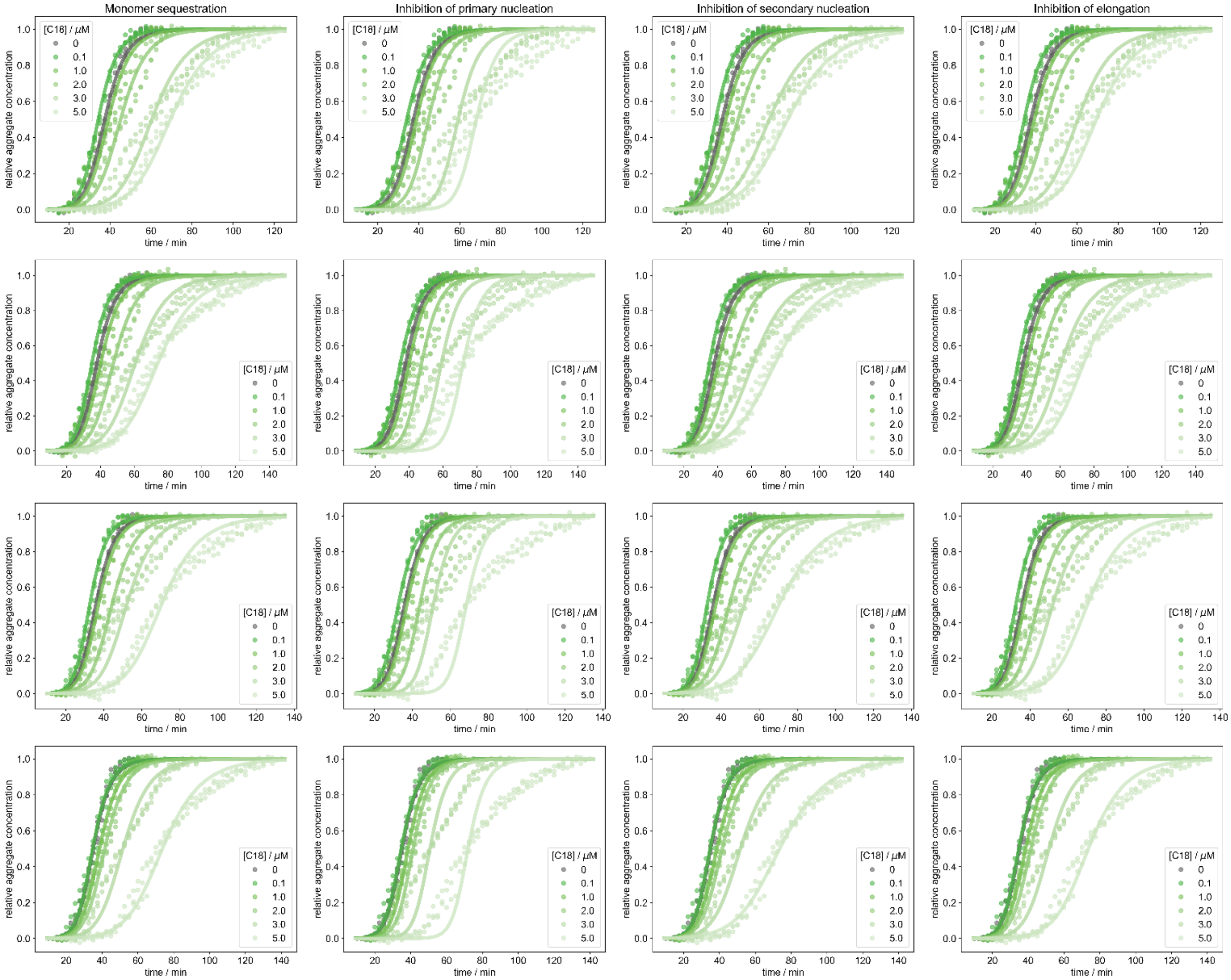


**Fig. S13.** Fitting results (solid lines) of the aggregation of 2.5 μM Aβ42 in the presence of **C18**, by means of the normalised kinetics measured in thioflavin T fluorescence assays. The fits are based on the kinetic parameters obtained from the unperturbed systems, allowing deviations in only the total monomer concentration, the rate constant of primary nucleation, secondary nucleation or elongation, respectively (columns left to right). Different rows show independent repeats of the experiment.

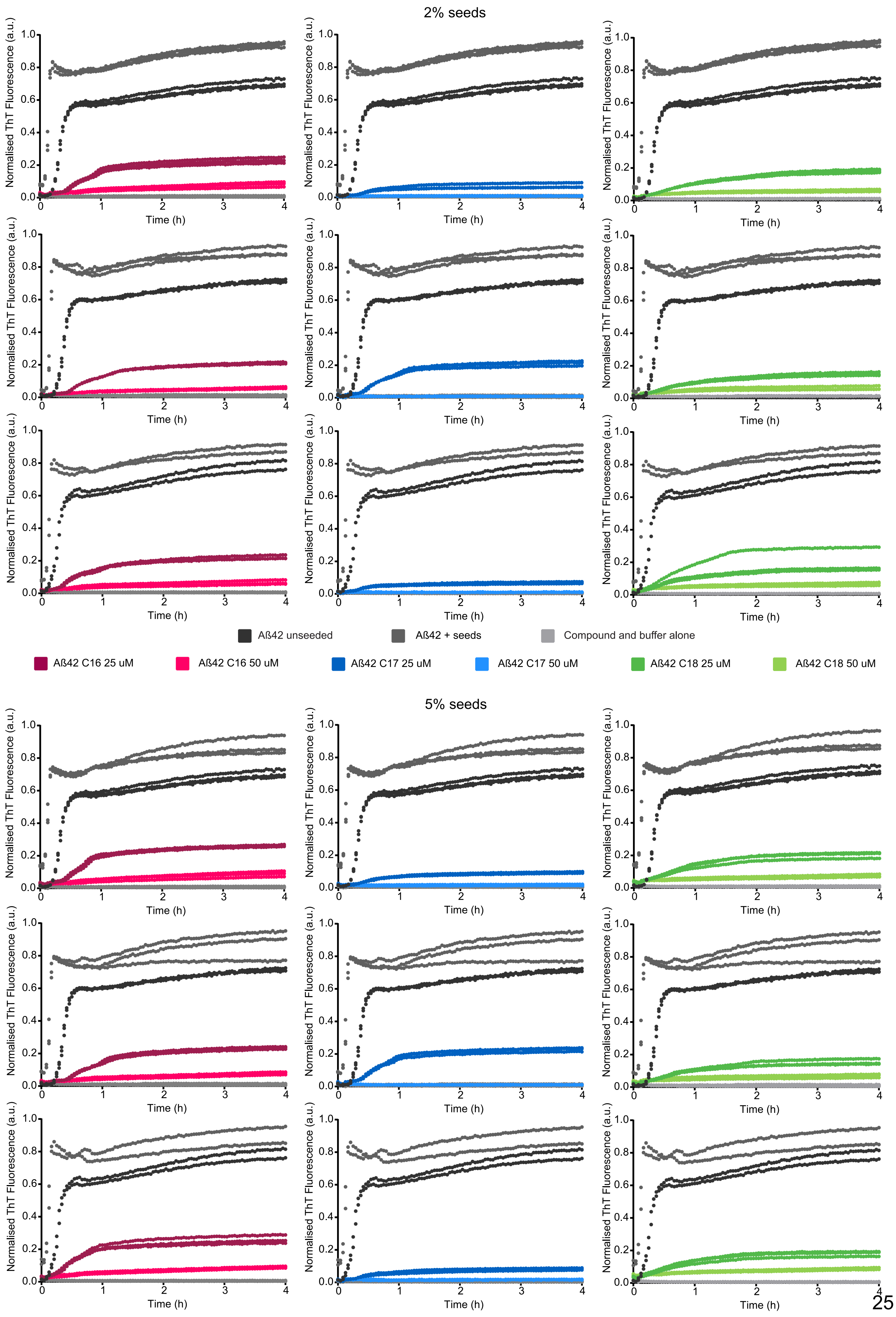
2% seeds
5% seeds
Normalised ThT Fluorescence (a.u.)
Time (h)
Aβ42 unseeded
Aβ42 + seeds
Compound and buffer alone
Aβ42 C16 25 uM
Aβ42 C16 50 uM
Aβ42 C17 25 uM
Aβ42 C17 50 uM
Aβ42 C18 25 uM
Aβ42 C18 50 uM

**Fig. S14.** Time-courses showing the effects of **C16**, **C17** and **C18** on seeded Aβ42 aggregation. 10 μM monomeric Aβ42 with 2% or 5% seeds in the absence or presence of either 1% DMSO or 25 or 50 μM of **C16** (pink), **C17** (blue) and **C18** (green). Samples were measured in triplicate under quiescent conditions at 37 °C. Data were corrected for time zero and normalised as a function of the maximum fluorescence intensity using GraphPad Prism.

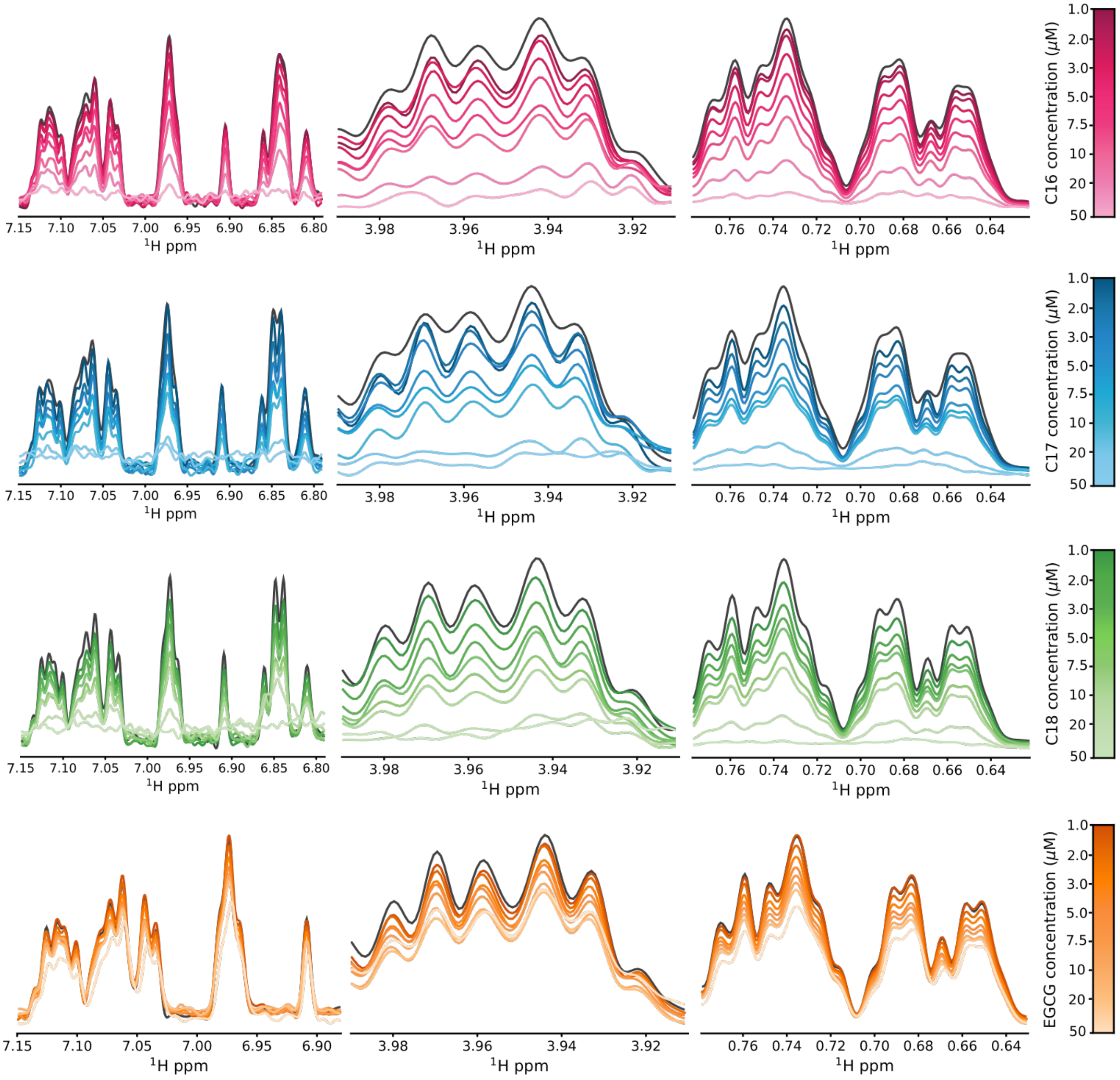


**Fig. S15.** One-dimensional $^1$H NMR spectra of Aβ42 with **C16** (pink), **C17** (blue), **C18** (green) and EGCG (orange). Unlabelled 10 μM Aβ42 (dark grey) was incubated with 1-50 uM compound and spectra recorded at 800 MHz at 4 °C. Three sections of the spectrum were analysed to determine the apparent $K_D$ for **C16**, **C17** and **C18**. *Left* N-H sidechain protons. *Middle* $H_\alpha$ protons. *Right* Methyl protons.

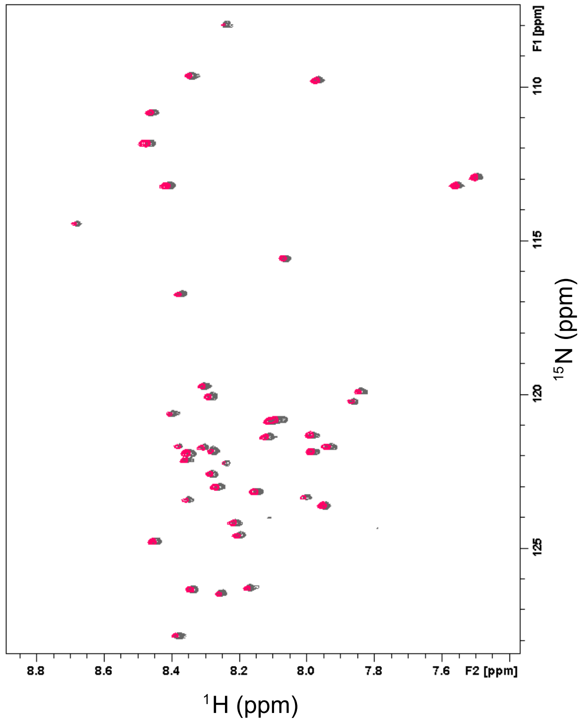


**Fig. S16.** Overlay of the $^{15}$N-$^{1}$H HSQC spectra of Aβ42 alone (grey) and Aβ42 with **C16** (pink). Spectra was recorded at 800 MHz at 4 °C.
